# Community venomics reveals intra-species variations in venom composition of a local population of *Vipera kaznakovi* in Northeastern Turkey

**DOI:** 10.1101/503276

**Authors:** Daniel Petras, Benjamin-Florian Hempel, Bayram Göçmen, Mert Karis, Gareth Whiteley, Simon C. Wagstaff, Paul Heiss, Nicholas R. Casewell, Ayse Nalbantsoy, Roderich D. Süssmuth

**Author notes:** These authors contributed equally to this study (Daniel Petras, Benjamin-Florian Hempel).

## Abstract

We report on the variable venom composition of a population of the Caucasus viper (*Vipera kaznakovi*) in Northeastern Turkey. We applied a combination of venom gland transcriptomics, as well as de-complexing bottom-up and top-down venomics, enabling the comparison of the venom proteomes from multiple individuals. In total, we identified peptides and proteins from 15 toxin families, including snake venom metalloproteinases (svMP; 37.8%), phospholipases A_2_ (PLA_2_; 19.0%), snake venom serine proteinases (svSP; 11.5%), C-type lectins (CTL; 6.9%) and cysteine-rich secretory proteins (CRISP; 5.0%), in addition to several low abundant toxin families. Furthermore, we identified intra-species variations of the *V. kaznakovi* venom composition, and find these were mainly driven by the age of the animals, with lower svSP abundance in juveniles. On a proteoform level, several small molecular weight toxins between 5 and 8 kDa in size, as well as PLA_2_s, drove the difference between juvenile and adult individuals. This study provides first insights into venom variability of *V. kaznakovi* and highlights the utility of intact mass profiling for a fast and detailed comparison of snake venoms of individuals from a community.

**Biological Significance:** Population level and ontogenetic venom variation (e.g. diet, habitat, sex or age) can cause a loss of antivenom efficacy against snake bites from wide ranging snake populations. The state of the art for the analysis of snake venoms are de-complexing bottom-up proteomics approaches. While useful, these have the significant drawback of being time-consuming and following costly protocols, and consequently are often applied to pooled venom samples. To overcome these shortcomings and to enable rapid and detailed profiling of large numbers of individual venom samples, we integrated an intact protein analysis workflow into a transcriptomics-guided bottom-up approach. The application of this workflow to snake individuals of a local population of *V. kaznakovi* revealed intra-species variations in venom composition, which are primarily explained by the age of the animals, and highlighted svSP abundance to be one of the molecular drivers for the compositional differences.

**Highlights:**

- First community venomic analysis of a local population of the Caucasian viper (*Vipera kaznakovi*).
- The venom gland transcriptome of *V. kaznakovi* identified 46 toxin genes relating to 15 venom toxin families.
- Bottom-up venomics revealed the identification of 25 proteins covering 7 toxin families mainly dominated by snake venom metalloproteinases (svMP).
- Community venomics by top-down mass profiling revealed ontogenetic shifts between juvenile and adult snakes.

## 1. Introduction

Venomics is considered an integrative approach, combining proteomics, transcriptomics and genomics to study venoms [1]. Although the term was initially used to describe the mass spectrometry-based proteomic characterization of venoms [2,3], genomic [4,5] or more commonly venom gland transcriptomic sequencing [6–14] have also been used to characterize venom compositions. These molecular approaches render an overview over venom composition by providing the nucleotide sequences of venom toxin-encoding genes (among others) and, in the case of transcriptomics, provide an estimation of their relative expression in the venom gland. Furthermore, (translated) protein sequence databases are crucial for the robust annotation of tandem mass spectra from proteomic analyses in peptide/protein spectrum matching (P*r*SM). A bibliographic search to the keyword “Snake venomics” in PubMed identified 147 hits between 2004 and 2018, which showed particularly in recent years a rapid expansion in the application of venomics approaches.

Initial proteomic analyses of snake venoms included the combination of multidimensional separation techniques (chromatographic and gel electrophoresis), N-terminal Edman degradation, and *de novo* sequencing by tandem mass spectrometry of tryptic peptides gathered by in-gel digestion of SDS-PAGE bands [2,15]. Since these initial studies, the proteomic characterization of snake venoms has become much more comprehensive due to technical advances in mass spectrometry and next generation nucleotide sequencing. Several complementary strategies were developed to unveil the venom proteomes of more than 100 snake species [16]. Most of these studies applied the so called ‘bottom-up’ proteomics whereby intact proteins are typically digested with trypsin before tandem mass spectrometry analysis. Many workflows perform venom decomplexation prior to the digestion either by liquid chromatography (LC) or gel electrophoresis, or a combination of both [17]. The direct, in-solution digestion, or so called ‘shotgun proteomics’, allows for a fast qualitative overview, but suffers from a less quantitative breakdown of snake venom composition [17,18]. For example, in shotgun experiments, the problem of protein inference often does not permit the differentiation of the numerous toxin isoforms present in venom [19]. Thus, the chromatographic or electrophoretic separation of venom samples greatly aids in differentiating between toxin isoforms (paralogs). In addition, decomplexing prior to trypsin digestion often does not allow for a clear identification of differential post-translational modified variants, so-called proteoforms [20].

A logical bypass of this problematic would be the omittance of the digestion step and the direct analysis of intact proteins by tandem mass spectrometry, called top-down proteomics. Recently top-down protein analysis has been applied alone or in combination with other venomics approaches to study the venom of King Cobra (*Ophiophagus hannah*) [21,22] the entire genus of mambas (*Dendroaspis* spp.) [23,24], the brown tree snake (*Boiga irregularis*) [6], the Okinawa Habu Pit Viper (*Protobothrops flavoviridis*) [25] and several viper species from Turkey [26–28]. In the case of viperid species, top-down analysis typically only reveals a partial characterization of the venom, as a number of the main toxin components, such as high molecular weight snake venom metalloproteinases (svMPs) (>30 kDa), are challenging to efficiently ionize by denaturing electrospray ionization (ESI) and might only provide few observable fragments in tandem MS [29]. A possible way to overcome difficulties in terms of ionization of high molecular weight proteins is the application of native ESI, as described by Melani *et al*. [22]. However native top-down mass spectrometry typically requires a special type of mass spectrometer with extended mass range and more extensive sample preparation, which makes this type of analysis more technically challenging. In most of the aforementioned studies, the top-down workflow was performed with a front-end LC-based sample decomplexation. This allows for the generation of MS1 mass profiles (XICs) of intact proteoforms. Typically, the MS1 information is accompanied by tandem MS (MS2) information acquired in data-dependent acquisition (DDA) mode. The MS2 fragment spectra are than matched to a translated transcriptome/genome database in order to identify the proteins. In the case that there are not enough MS2 fragment peaks of a particular proteoform, the intact molecular mass can still enable identification, especially if the intact mass can be associated to masses observed in complementary experiments, such as retention time, mass range of SDS-PAGE and/or bottom-up protein IDs of decomplexed bottom-up venomics [26]. The additional information gained through exact intact protein masses can be particularly informative to differentiate between isoforms or proteoforms. Furthermore, the simple sample preparation, high sensitivity and fast analysis time allows for a rapid comparison of the venom composition. The quantitative capabilities of top-down approaches [30–32] thereby offer great potential for comparative venomic studies of individuals. While most snake venom compositions reported so far [16] were performed on a single pool of venom sourced from different numbers of individuals, several studies have shown correlations between different ecological, geographical, genetic and/or developmental factors and the venom proteome, e.g. different diets [33–36], regional separation of populations [37–39], sex [40–42] or age [43–46]. In addition to better understand the heritability of venom toxins [47], and the evolutionary processes underpinning population level venom variations [48], venomics is an important approach to better understand regional and intraspecific variations in the venom composition of medically important snake species, which has considerable relevance for the development and clinical utility of snakebite therapies, known as antivenom [49,50].

Here, we applied a top-down venomics approach to demonstrate intraspecific venom variation in a local population of the medical relevant Caucasian viper (*Vipera kaznakovi*). The Caucasian viper is a subtropical, medium-sized, viper species with a distribution range mainly at the Caucasian Black Sea coast in the Artvin and Rize province of Turkey. *V. kaznakovi* feed predominately on small vertebrates (mice, lizards etc.) and also insects [51]. In a previous shotgun proteomics study of this species, Utkin and coworkers described the venom of *V. kaznakovi* to be composed of phospholipase A_2_ (PLA_2_), svMP, snake venom serine proteases (svSP), Cysteine-rich secretory proteins (CRISP), C-type lectins (CTL), L-amino acid oxidase (LAAO), vascular endothelial growth factor (VEGF), disintegrins (Dis), phospholipase B (PLB), nerve growth factors (NGF), as well as other a number of other proteins of lower abundance [52]. In this study, we pursue a more in-depth approach to characterizing the venom of this species. We use a combination of venom gland transcriptomics, decomplexing bottom-up proteomics and comparative top-down proteomics to broadly characterize the venom composition of this species, and to also investigate intraspecific variation in toxins on the level of the individual snakes.

## 2. Material & Methods

### 2.1. Sampling

Venom samples of *V. kaznakovi* were collected from 6 adult (2 female, 4 male) and 3 juvenile specimens (unknown sex). All specimens were captured in late June 2015 in their natural habitat and released back into their natural environment after venom extraction. The *V. kaznakovi* individuals were collected in Artvin province in Turkey near the Georgian border, with 6 individuals sampled from Hopa district, 2 individuals from Borçka district and 1 specimen in the Arhavi district. An additional female individual found in Borçka district was collected for venom gland dissection for transcriptomics analysis. Ethical permission (Ege University Animal Experiments Ethics Committee, 2013#049) and special permission (2015#124662) for the sampling of wild-caught *V. kaznakovi* were received from the Republic of Turkey, Ministry of Forestry and Water Affairs.

### 2.2. Sample storage and preparation

Crude *V. kaznakovi* venom was extracted by using a parafilm-covered laboratory beaker without exerting pressure on the venom glands. Venom samples were centrifuged at 2000 × g for 10 min at 4 °C to remove cell debris. Supernatants were immediately frozen at −80 °C, lyophilized and samples were stored at 4 °C until use.

### 2.3. Determination of lethal dose (LD_50_)

Lethal potency (LD_50_) of venoms to mice (milligrams of dry weight per kg) was determined by an up- and down method as recommended by the Organization for Economic Cooperation and Development (OECD) guidelines (Test No. 425) [53,54]. Groups of five mice (n = 15; age, 8 to 10 weeks; female 8 and male 7 individuals) were used per venom dose. Various venom concentrations (5, 2 and 1 mg/kg, milligrams of protein per kg calculated from dry weight venom by Bradford assay) were diluted in ultrapure water to a final volume of 100 μL and injected by intraperitoneal (IP) routes. Control mice (n = 5; female 2 and male 3 individuals) received a single IP injection of sterile saline (0.9%, 0.1 mL). All assays and procedures involving animals strictly followed the ethical principles in animal research adopted by the Swiss Academy of Medical Sciences [55]. Additionally, they were approved by a local ethics committee (2013#049). The mortality was recorded 24 h after injection. The median lethal dose was determined by a nonlinear regression fitting procedure (GraphPad Prism 5., Version 5.01, Inc., San Diego, CA, USA).

### 2.4. RNA isolation and purification

Venom glands were dissected from a wild caught adult female specimen of *V. kaznakovi* in Kanlıdere, Hopa district (Artvin province) and processed as previously described [9,24]. Briefly, immediately following euthanasia, venom glands were dissected and were immediately flash frozen in liquid nitrogen and stored cryogenically prior to RNA extraction. Venom glands were next homogenized under liquid nitrogen and total RNA extracted using a TRIzol Plus RNA purification kit (Invitrogen), DNAse treated with the PureLink DNase set (Invitrogen) and poly(A) selected using the Dynabeads mRNA DIRECT purification kit (Life Technologies), as previously detailed [9,24].

### 2.5. RNA sequencing, assembly and annotation

RNA-Seq was performed as previously described [9,24]. The RNA-Seq library was prepared from 50 ng of enriched RNA material using the ScriptSeq v2 RNA-Seq Library Preparation Kit (epicenter, Madison, WI, USA), following 12 cycles of amplification. The resulting sequencing library was purified using AMPure XP beads (Agencourt, Brea, CA, USA), quantified using the Qubit dsDNA HS Assay Kit (Life Technologies), before the size distribution was assessed using a Bioanalyser (Agilent). The library was then multiplexed and sequenced (alongside other sequencing libraries not reported in this study) on a single lane of an Illumina MiSeq, housed at the Centre for Genomic Research, Liverpool, UK. The *V. kaznakovi* library amounted to 1/6^th^ of the total sequencing lane. The ensuing read data was quality processed by (i) removing the presence of any adapter sequences using Cutadapt (https://code.google.com/p/cutadapt/) and (ii) trimming low quality bases using Sickle (https://github.com/najoshi/sickle). Reads were trimmed if bases at the 3’ end matched the adapter sequence for 3 bp or more, and further trimmed with a minimum window quality score of 20. After trimming, reads shorter than 10 bp were removed.

For sequence assembly we used VTBuilder, a *de novo* transcriptome assembly program previously designed and validated for constructing snake venom gland transcriptomes [56]. Paired-end read data was entered into VTBuilder and executed with the following parameters: min. input read length 150 bp; min. output transcript length 300 bp; min. isoform similarity 96%. Assembled contigs were annotated with the BLAST2GO Pro v3 [57] using the blastx-fast algorithm with a significance threshold of 1e^−5^, to provide BLAST annotations (max 20 hits) against NCBI’s non redundant (NR) protein database (41 volumes; Nov 2015) followed by mapping to gene ontology terms, and Interpro domain annotation using default parameters. Following generic annotation, venom toxins were initially identified based on their BLAST similarity to sequences previously identified in the literature or in molecular databases as snake venom toxins, and then manually curated for validation.

### 2.6. Venom proteomics (bottom-up)

The crude venom (1 mg) was dissolved to a final concentration of 10 mg/ml in aqueous 3% (v/v) acetonitrile (ACN) with 1% (v/v) formic acid (FA) and centrifuged at 16,000 *g* for 5 min to spin down insoluble content. The supernatant was loaded onto a semi-preparative reversed-phase HPLC with a Supelco Discovery BIO wide Pore C18–3 column (4.6 × 150 mm, 3 μm particle size) using an Agilent 1260 Low Pressure Gradient System (Agilent, Waldbronn, Germany). The column was operated with a flow rate of 1 mL/min and performed with ultrapure water (solution A) and ACN (solution B), both including 0.1% (v/v) FA. A standard separation gradient was used with solution A and solution B, starting isocratically (5% B) for 5 min, followed by linear gradients of 5–40% B for 95 min and 40–70% for 20 min, then 70% B for 10 min, and finally re-equilibration at 5% B for 10 min. Peak detection was performed at *λ* = 214 nm using a diode array detector (DAD). After the chromatographic separation of the crude venom, the collected and vacuum-dried peak fractions were submitted to a SDS-PAGE gel (12% polyacrylamide). Subsequently, the coomassie-stained bands were excised, and submitted to in-gel trypsin digestion, reduced with fresh dithiothreitol (100 mM DTT in 100 mM ammonium hydrogencarbonate, pH 8.3, for 30 min at 56 °C) and alkylated with iodoacetamide (55 mM IAC in 100 mM ammonium hydrogencarbonate, pH 8.3, for 30 min at 25 °C in the dark). The resulting peptides were then extracted with 100 μL aqueous 30% (v/v) ACN just as 5% (v/v) FA for 15 min at 37 °C. The supernatant was vacuum-dried (Thermo speedvac, Bremen, Germany), redissolved in 20 μL aqueous 3% (v/v) ACN with 1% (v/v) FA and submitted to LC-MS/MS analysis.

The bottom-up analysis were performed with an Orbitrap XL mass spectrometer (Thermo, Bremen, Germany) via an Agilent 1260 HPLC system (Agilent Technologies, Waldbronn, Germany) using a reversed-phase Grace Vydac 218MSC18 (2.1 × 150 mm, 5 μm particle size) column. The pre-chromatographic separation was performed with the following settings: After an isocratic equilibration (5% B) for 1 min, the peptides were eluted with a linear gradient of 5–40% B for 10 min, 40–99% B in 3 min, held at 99% B for 3 min and re-equilibrated in 5% B for 3 min.

### 2.7. Community venom profiling (top-down)

The top-down MS analysis was performed by dissolving the crude venoms in ultrapure water containing formic acid (FA, 1%) to a final concentration of 10 mg/mL, and centrifuged at 20,000 × g for 5 min. Aliquots of 10 μL dissolved venom samples were submitted to reverse-phase (RP) HPLC-high-resolution (HR)-MS analyses. RP-HPLC-HR-MS experiments were performed on an Agilent 1260 HPLC system (Agilent, Waldbronn, Germany) coupled to an Orbitrap LTQ XL mass spectrometer (Thermo, Bremen, Germany). RP-HPLC separation was performed on a Supelco Discovery Biowide C18 column (300 Å pore size, 2 × 150 mm column size, 3 μm particle size). The flow rate was set to 0.3 mL/min and the column was eluted with a gradient of 0.1% FA in water (solution A) and 0.1% FA in ACN (solution B): 5% B for 5 min, followed by 5–40% B for 95 min, and 40–70% for 20 min. Finally, the gradient was held isocratic with 70% B for 10 min and re-equilibrated at 5% B for 10 min. ESI settings were: 11 L/min sheath gas; 35 L/min auxiliary gas; spray voltage, 4.8 kV; capillary voltage, 63 V; tube lens voltage, 135 V; and capillary temperature, 330 °C. MS/MS spectra were obtained in data-dependent acquisition (DDA) mode. FTMS measurements were performed with 1 μ scans and 1000 ms maximal fill time. AGC targets were set to 10^6^ for full scans and to 3 × 10^5^ for MS/MS scans, and the survey scan as well as both data dependent MS/MS scans were performed with a mass resolution (R) of 100,000 (at *m/z* 400). For MS/MS the two most abundant ions of the survey scan with known charge were selected. Normalized CID energy was set to 30% for the first, and 35% for the second, MS/MS event of each duty cycle. The default charge state was set to z = 6, and the activation time to 30 ms. Additional HCD experiments were performed with 35% normalized collision energy, 30 ms activation time and z = 5 default charge state. The mass window for precursor ion selection was set to 2 or 6 *m/z*. A window of 3 *m/z* was set for dynamic exclusion of up to 50 precursor ions with a repeat of 1 within 10 s for the next 20 s.

### 2.8. Bioinformatic analysis

The LC-MS/MS data files (.raw) obtained from the in-gel digestion were converted to mascot generic format (.mgf) files via MSConvert GUI of the ProteoWizard package (http://proteowizard.sourceforge.net; version 3.0.10328) and annotated by DeNovo GUI [58] (version 1.14.5) with a mass accuracy of 10 ppm for precursor mass and 0.2 m/z for fragment peaks. A fixed modification carbamidomethyl cysteine (C +57.02 Da) was selected. Resulting sequence tags were examined manually and searched against the non-redundant Viperidae protein database (taxid: 8689) using the basic local alignment search tool (BLAST) [59].

For peptide spectrum matching, the SearchGUI software tool was used with XTandem! As the search engine [60]. The MS2 spectra were searched against the non-redundant Viperidae protein NCBI (taxid: 8689, 3rd Nov 2017, 1727 sequences), our in-house *Vipera kaznakovi* toxin sequence database (translated from our venom gland transcriptomic analyses; 46 toxin sequences) and a set of proteins found as common contaminants (CRAP, 116 sequences), containing in total 1889 sequences. Mass accuracy was set to 10 ppm for the precursor mass and 0.2 m/z for the MS2 level. Alkylation of Cys was set as fixed modification and acetylation of the N-terminus, of Lys as well as oxidation of Met were allowed as variable modifications. A false discovery rate (FDR) was estimated through a target-decoy approach and a cut-off of 1% was applied. All PSMs were validated manually and at least 2 PSMs were required for a protein ID to be considered.

For the top-down data analysis, the .raw data were converted to .mzXML files using MSconvert of the ProteoWizard package (http://proteowizard.sourceforge.net; version 3.0.10328), and multiple charged spectra were deconvoluted using the XTRACT algorithm of the Xcalibur Qual Browser version 2.2 (Thermo, Bremen, Germany). For isotopically unresolved spectra, charge distribution deconvolution was performed using the software tool <u>mag</u>ic <u>tran</u>sformer (MagTran).

### 2.9. Multivariable statistics

Principal component analysis (PCoA), using the relative percentages of the major toxin families as well as different proteoforms as a variable, was applied to explain determinants of compositional variation among venoms. PCoA was performed in R (R Foundation for Statistical Computing, 2016) with the extension Graphic Package rgl, available from https://www.R-Project.org.

### 2.10. Data sharing

Mass spectrometry proteomics data (.mgf, .raw and results files and search database) have been deposited to ProteomeXchange [61] with the ID PXD010857 via the MassIVE partner repository under project name “Venom proteomics of *Vipera kaznakovi*” and massive ID MSV000082845.

Raw sequencing reads and the assembled contigs generated for the venom gland transcriptome (.fastq and .fasta, respectively) have been deposited in the NCBI sequence read archive (SRA) under accession SRR8198764 and linked to the BioProject identifier PRJNA505487.

## 3. Results and Discussion

### 3.1. Field work and venom toxicity

The medium-sized Caucasian viper (*Vipera kaznakovi*) mainly inhabits the forested slopes of mountain peaks with a distribution range from the Caucasian Black Sea coast provinces of northeastern Turkey, through Georgia to Russia (**Figure 1**). *V. kaznakovi* feeds predominately on small vertebrates (mice, lizards, etc.) or insects, and a specific characteristic of this species is the complete black coloration with elements of orange to red zigzag-looking strip on the upper side of the body (**Figure 1**).

**Figure 1.**
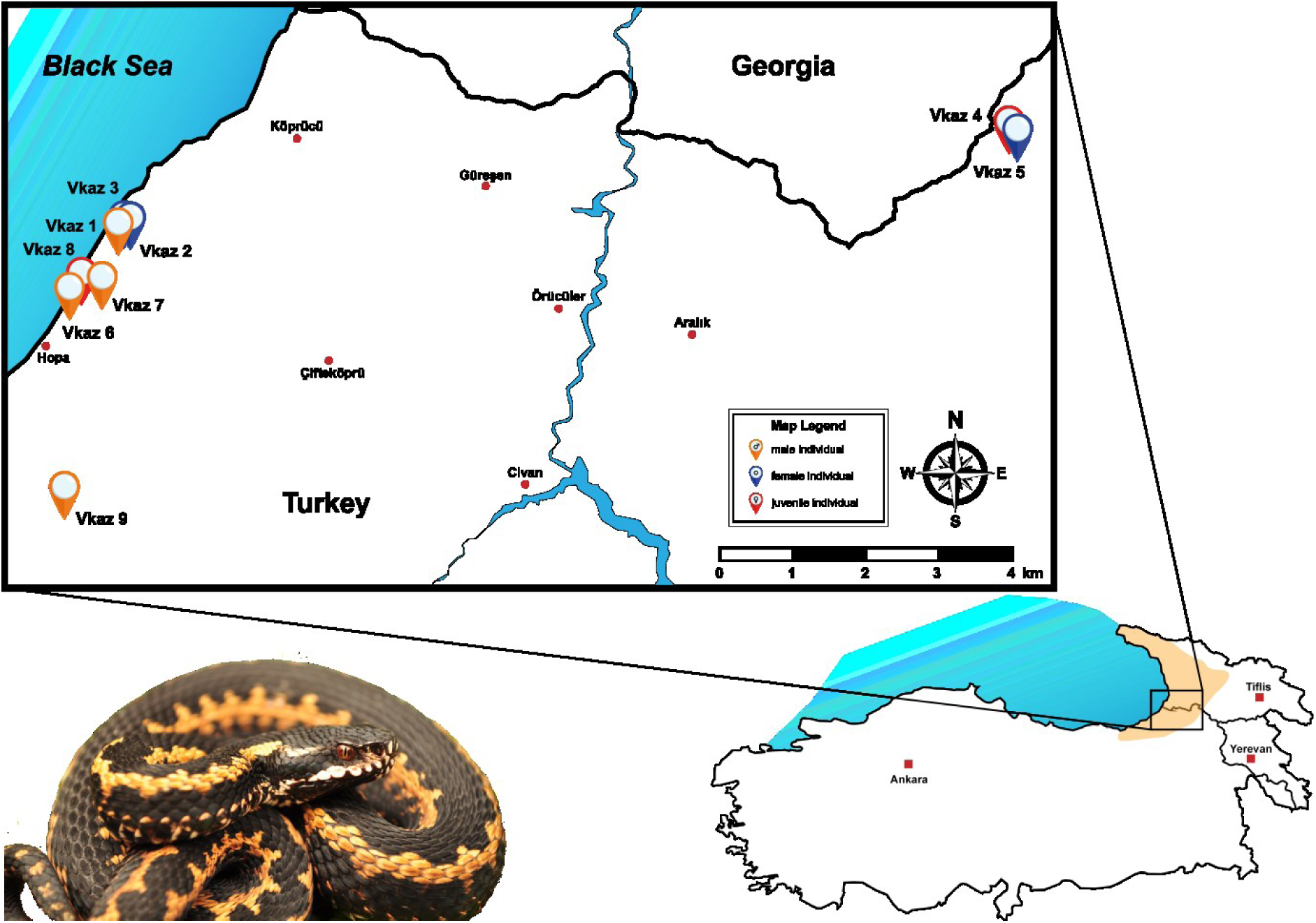
Geographical distribution and sampling localities of *Vipera kaznakovi*. The distribution area of the Caucasian Viper (*Vipera kaznakovi*, genus *Viperidae*) is highlighted on the map in the lower right corner and adapted from Geniez *et al*. [83]. The locations and sex/age of the collected individuals are marked on the map (orange – adult male, red – adult female, blue - juvenile).

During our fieldwork in June 2015 we collected nine *V. kaznakovi* individuals (6 adults and 3 juveniles) in their natural habitat, whose venom was extracted by using a parafilm-covered laboratory beaker before the snakes were released back into their natural environment. The different *V. kaznakovi* individuals were found in Hopa (6 spec.), Borçka (2 spec.) and Arhavi (1 spec.) districts of the Artvin province (**Figure 1**). The LD_50_ mean values of venom pooled from all collected *V. kaznakovi* individuals was assessed by the intraperitoneal (IP) route using a random sample survey of five swiss mice for three venom dose (5, 2 and 1 mg/kg), which is summarized in supplemental table 1. The LD_50_ mean value obtained for the pooled *V. kaznakovi* venom was calculated as ~2.6 mg/kg and can categorized to have slightly weaker toxicity in this model, compared to other related viper species (0.9–1.99 mg/kg) [62–65].

**Table 1.**
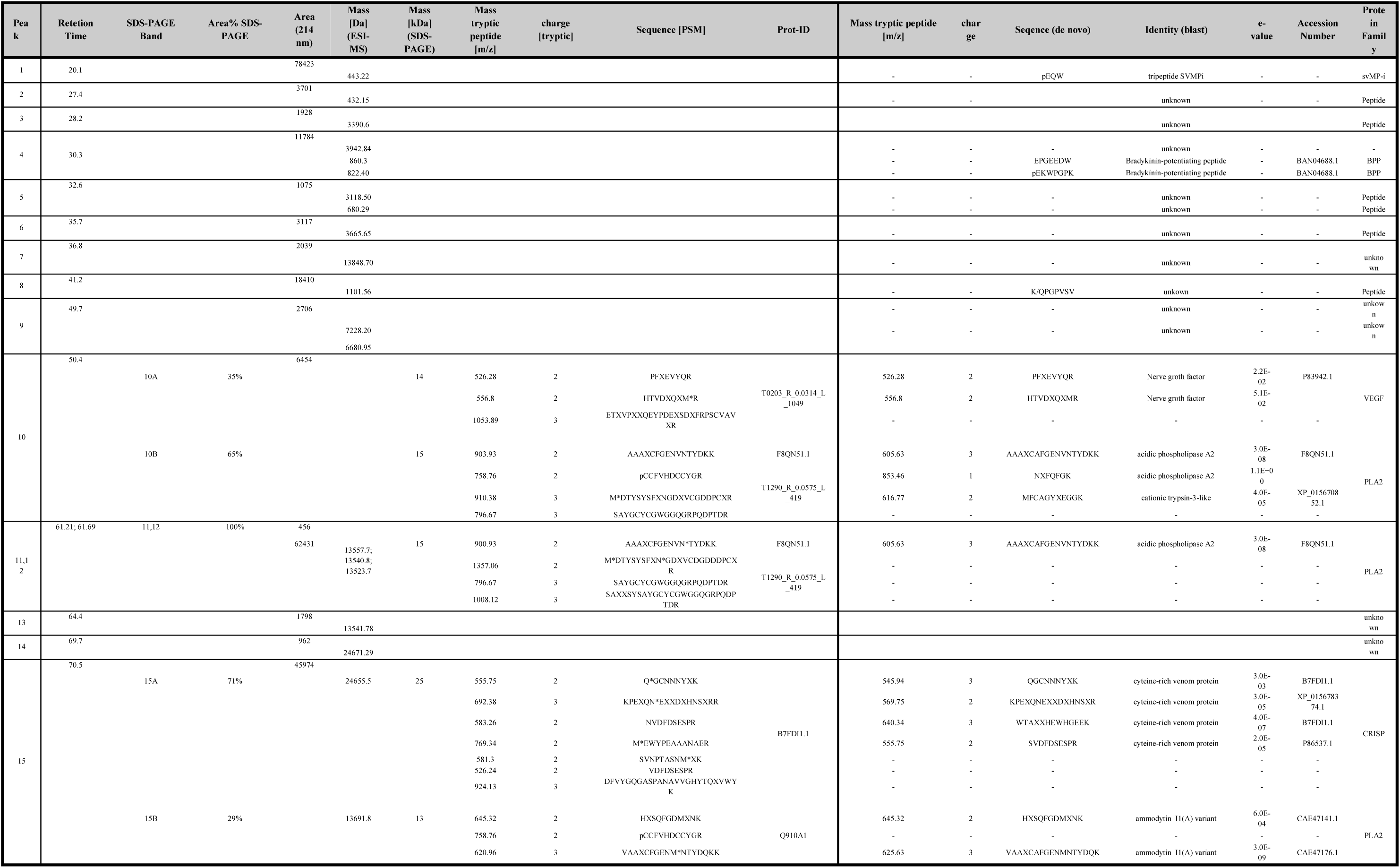
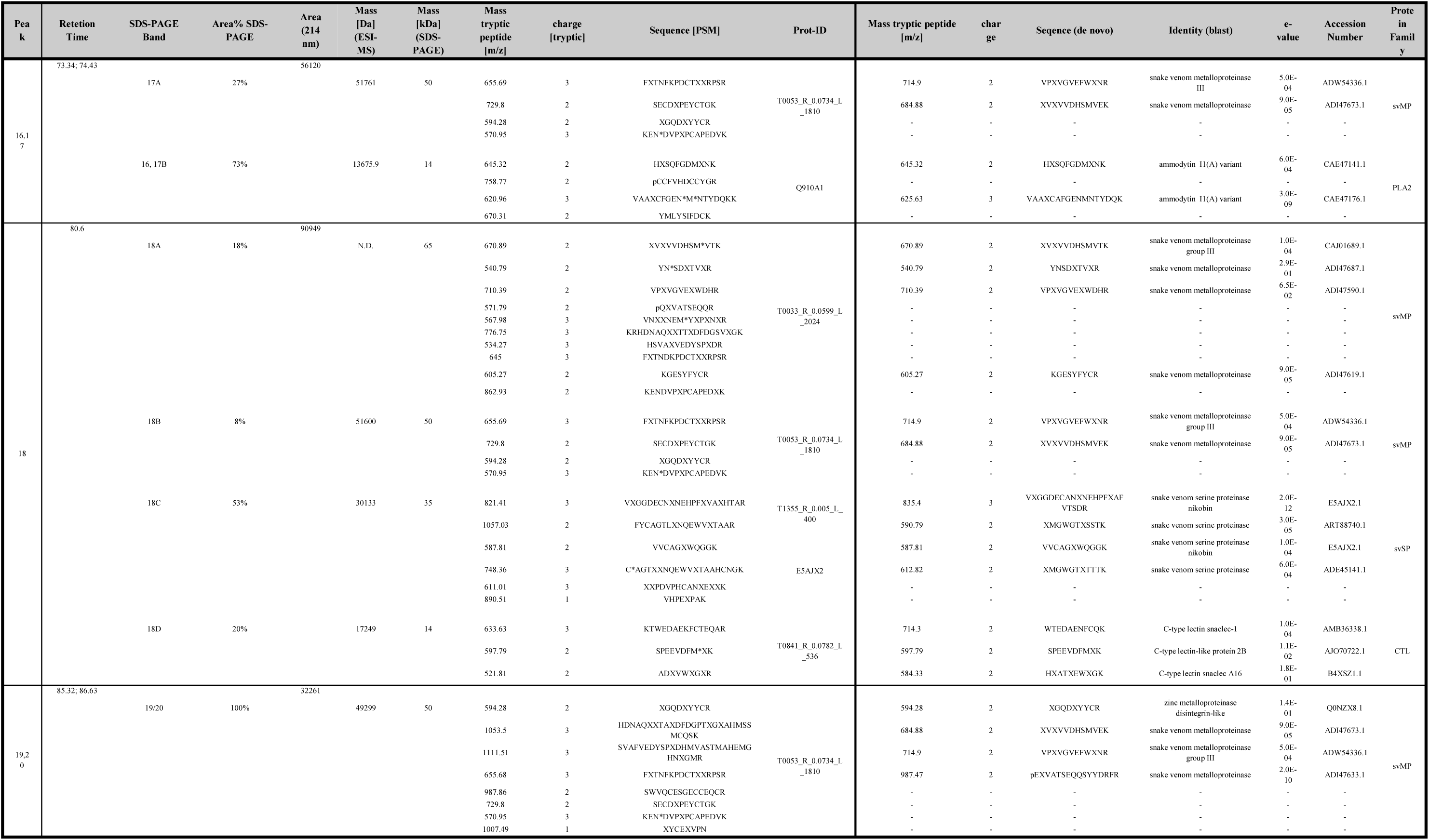
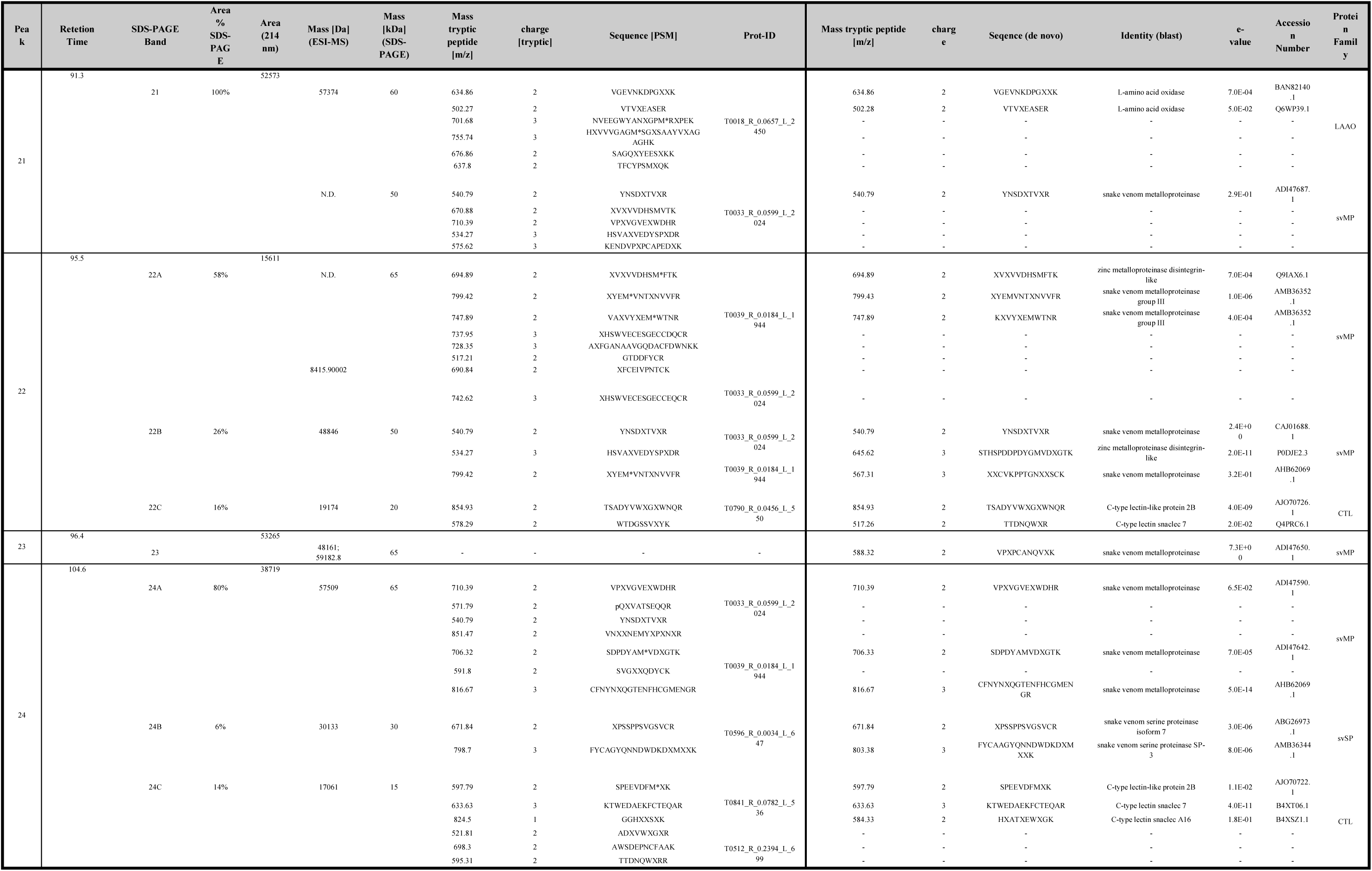
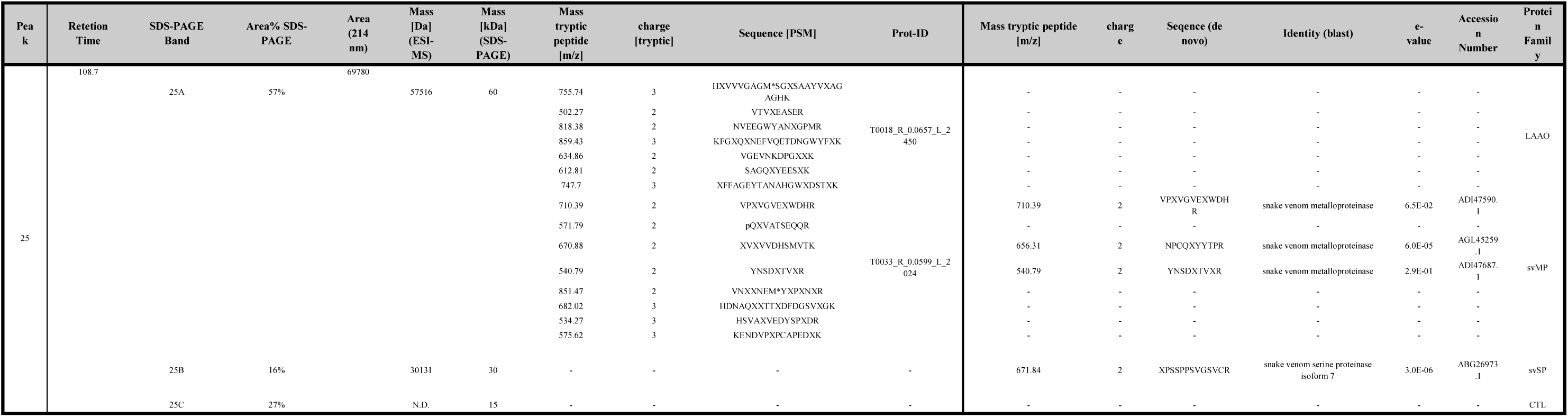
Venom Protein Identifications from *Vipera kaznakovi*. The table shows all protein identification of HPLC fractions (Fig. 3) by LC-MS and LC-MS/MS analysis from pooled venom. Peak numbering corresponds to the UV and MS chromatograms. Sequence tags were obtained by analysis of tryptic peptides by MS/MS de novo sequencing and/or peptide spectrum matching. Molecular weights of intact proteins were determined by SDS-PAGE and intact mass profiling (LC-MS).

### 3.2. Venom gland transcriptomics

The *V. kaznakovi* venom gland transcriptome resulted in 1,742 assembled contigs, of which 46 exhibited gene annotations relating to 15 venom toxin families previously described in the literature (**Figure 2**). The majority of these contigs (33) encode genes, expressing toxin isoforms relating to four multi-locus gene families, namely the svMPs, CTLs, svSPs and PLA_2_s (**Figure 2**). Moreover, these four toxin families also exhibited the highest expression levels of the toxin families identified; in combination accounting for >78% of all toxin expressions (**Figure 2**). These findings are consistent with many prior studies of viperid venom gland transcriptomes [10,12,49,66,67].

**Figure 2.**
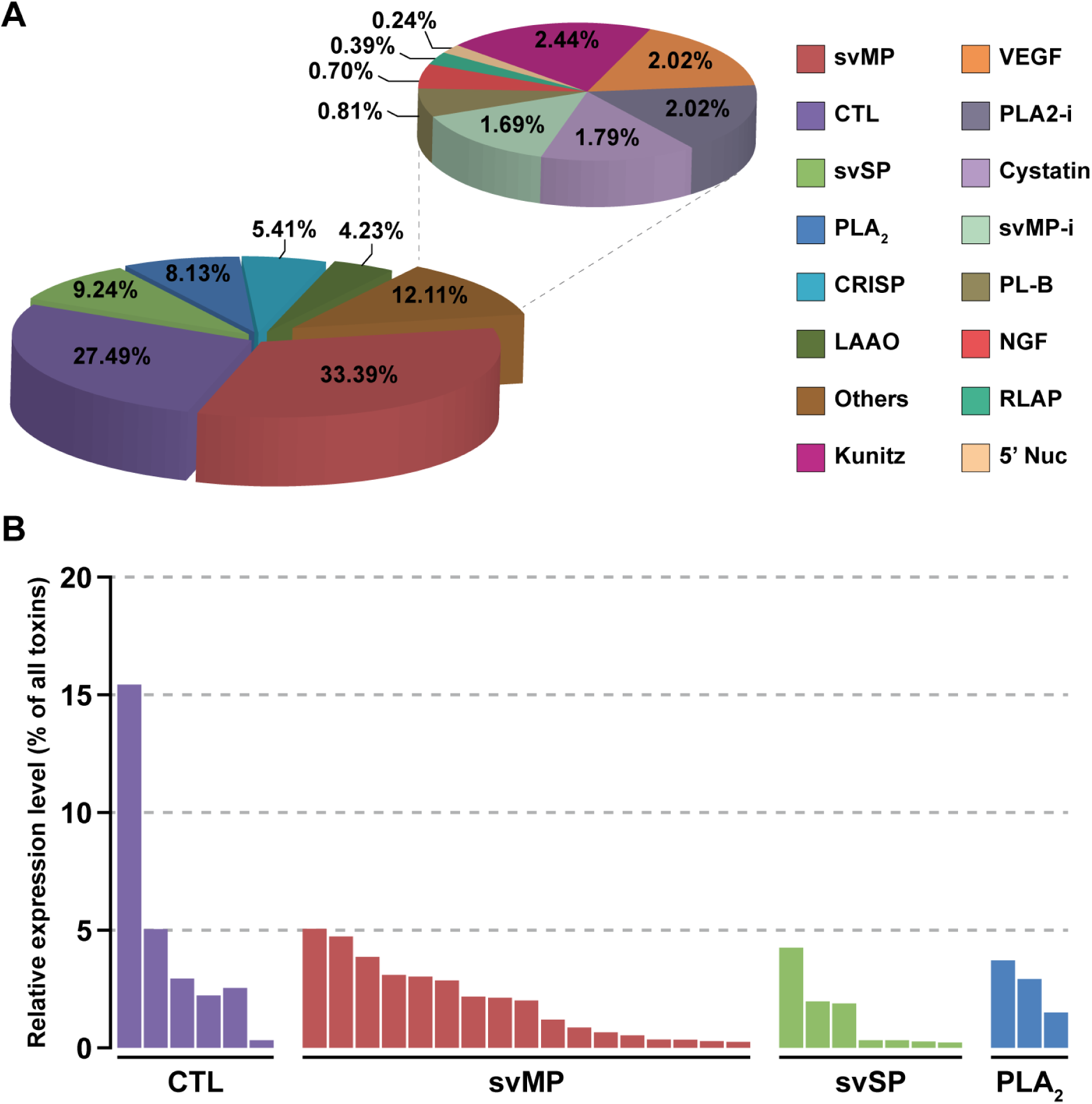
The relative expression levels of toxin families identified in the *Vipera kaznakovi* venom gland transcriptome. **A** The left pie chart shows the relative expression levels of the major toxin families, each of which accounts for greater than 4% of all toxins encoded in the venom gland. The right pie chart shows the relative expression levels of the remaining toxin families, which in combination account for 12.11% of all toxins encoded in the venom gland (“others”). Percentage values on both charts reflect the expression level of each toxin family as a percentage of the total expression of all identified toxin families. **B** The relative expression levels of individual contigs encoded by the most abundantly expressed toxin families (CTL, svMP, svSP and PLA_2_). Key: svMP – snake venom metalloproteinase; CTL – C-type lectin; svSP – snake venom serine protease; PLA_2_ – phospholipase A_2_; CRISP – cysteine-rich secretory protein; LAAO – L-amino acid oxidase; kunitz – kunitz-type inhibitors; VEGF – vascular endothelial growth factor; PLA2-i – PLA_2_ inhibitors, SVMP-i – SVMP inhibitors PLB – phospholipase B; NGF – nerve growth factor; RLAP – renin-like aspartic proteases; 5’ Nuc – 5’ nucleotidase.

The svMPs were the most abundantly expressed of the toxin families, accounting for 33.4% of the total toxin expression, and were encoded by 17 contigs (**Figure 2**). However, these contig numbers are likely to be an overestimation of the total number of expressed svMP genes found in the *V. kaznakovi* venom gland, as six of these contigs were incomplete and non-overlapping in terms of their nucleotide sequence, and therefore likely reflect a degree of low transcriptome coverage and/or under-assembly. Of those contigs that we were able to identify to svMP class level (e.g. P-I, P-II or P-III [68,69]), ten exhibited structural domains unique to P-III svMPs, one to P-II svMPs and one to a short coding disintegrin. Interestingly, the svMP contig that exhibited the highest expression level encoded for the sole P-II svMP (5.1% of all venom toxins), whereas the short coding disintegrin, which exhibited 98% identity to the platelet aggregation inhibitor lebein-1-alpha from *Macrovipera lebetina* [70], was more moderately expressed (2.1%). Interestingly, we found no evidence for the representation of the P-I class of svMPs in the *V. kaznakovi* venom gland transcriptome.

The CTLs were the next most abundant toxin family, with six contigs representing 27.5% of all toxin gene expression (**Figure 2**). Interestingly, one of these CTLs, which exhibits the closest similarity to snaclec-7 from *Vipera ammodtyes* venom (GenBank: APB93444.1), was by far the most abundantly expressed toxin identified in the venom gland transcriptome (15.4% of all toxins) (**Figure 2**). We identified lower expression levels for the multi-locus svSP and PLA_2_ toxin families, which accounted for 9.2% and 8.1% of the toxins, expressed in the venom gland transcriptome respectively, and were encoded by seven and three contigs (**Figure 2**). Of the remaining toxin families identified, only two exhibited expression levels >3% of the total toxin expression; CRISPs were encoded by two contigs amounting to 5.4% of total toxin expression, and LAAO by a single contig representing 4.23% (**Figure 2**). The remaining nine, lowly expressed, toxin families identified in the venom gland transcriptome are displayed in Figure 2, and combined amounted to 12.1% of total toxin expression.

### 3.3 Decomplexed proteomics of pooled venom

To broadly characterize the venom composition of *V. kaznakovi*, in an initial experiment, we performed bottom-up analysis of pooled venom by reversed phase-HPLC separation (**Figure 3A**) and direct online intact mass analysis by ESI-HR-MS (Figure **3B**). The prominent bands of the subsequent separation by SDS-PAGE (**Figure 3C**) were excised followed by trypsin in-gel digestion and LC-MS/MS analysis. During the first analysis we did not have a species-specific transcriptome database available, hence the spectra were analyzed by *de novo* sequencing. The resulting sequence tags were searched against the NCBI non-redundant viperid protein database using BLAST [59]. The 57 sequence tags resulted in the identification of 25 proteins covering 7 toxin families (**Table 1**), namely svMP, PLA_2_, svSP, CTL, CRISP, VEGF and LAAO.

**Figure 3.**
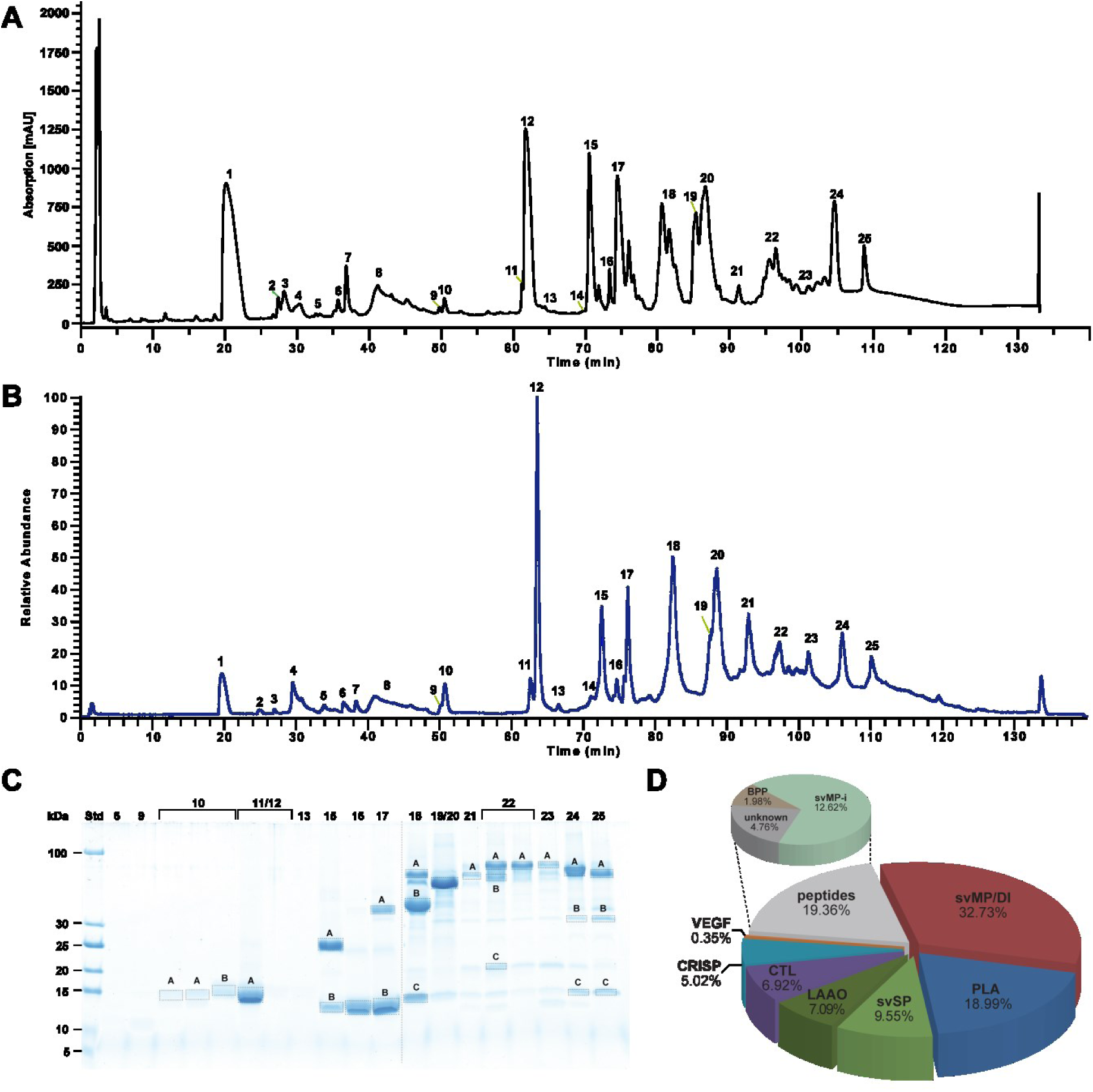
Bottom-up snake venomics of *Vipera kaznakovi*. **A** Venom separation of *V. kaznakovi* was performed by a Supelco Discovery BIO wide Pore C18–3 RP-HPLC column and UV absorbance measured at *λ* = 214 nm. **B** Total ion current (TIC) profile of crude *V. kaznakovi* venom. The peak nomenclature is based on the chromatogram fractions. **C** The RP-HPLC fractions (indicated above the lane) of the *V. kaznakovi* venom was analysed by SDS-PAGE under reducing conditions (Coomassie staining). Alphabetically marked bands per line were excised for subsequent tryptic in-gel digestion. **D** The relative occurrence of different toxin families of *V. kaznakovi* is represented by the pie chart. Identification of snake venom metalloproteinase (svMP, red), phospholipases A_2_ (PLA_2_, blue), snake venom serine proteinase (svSP, green), C-type lectin like proteins (CTL, purple), cysteine rich secretory proteins (CRISP, light blue), bradykinin-potentiating peptides (BPP, light brown), vascular endothelial growth factors (VEGF-F, red), unknown proteins (n/a, black) and peptides (grey). The *de novo* identified peptides are listed in supplemental table 1.

*De novo* sequencing of MS/MS spectra of native small peptides (peaks 1–9) resulted in four additional sequence tags and the identification of a svMP inhibitor (svMP-i) and two bradykinin potentiating peptides (BPP). When we obtained the assembled transcriptome data, we re-analyzed the MS/MS data from the tryptic peptides by peptide spectrum matching (PSM) using the translated protein sequences of the transcriptome as well as the NCBI viperidae protein database. PSM resulted in 114 peptide matches in total, which doubled the number of annotated spectra in comparison to the *de novo* annotation. The analysis revealed the same seven major toxin families as identified by the tryptic *de novo* tags, but showed with 29 identified proteins (compared to 25 by the above approach) a slight improvement. Not surprisingly, most of the peptide matches were from the transcriptome derived sequences and only six protein IDs came from other viperid sequences from the NCBI database. Relative quantification through integration of the UV-HPLC peaks and densitometric analysis of the SDS-gels revealed that the most abundant toxin families were svMP (37.7%), followed by PLA_2_ (19.0%), svSP (9.6%), LAAO (7.1%), CTL (6.9%), CRISP (5.0%), and VEGF (0.3%). In the small molecular mass range < 2kDa, SVMP-i contributed 12.6%, BPP 2.0%, and unknown peptides 4.0% to the overall venom composition (**Figure 3D**).

Comparing the abundance of venom toxins (**Figure 3D**) with transcriptomic predictions of expression (**Figure 2A**), we observed an overall positive correlation, but we noted some major differences, particularly relating to the CTLs: transcriptomic expression levels showed CTLs to be the second most abundant toxin family (27.5% of all toxin contigs) while proteomic analysis shows a much lower occurrence (6.9%). Interestingly, some of the molecular masses observed for CTLs (~20 kDa) during SDS-PAGE did not correspond to the expected molecular mass derived from the transcriptome sequence. As reported in other studies, we assume that some of the observed CTLs are hetero-dimers [71]. SvMPs showed highly consistent profiles, as both the most abundantly expressed (33.4%) and translated (32.7%) toxin family. Similarly, the svSPs (9.2%) and CRISPs (5.4%) exhibited transcription levels highly comparable to their relative protein abundance in venom (9.6% and 5.02%). A lower transcription level was shown for PLA_2_ (8.1%) in contrast to the two times higher protein level (19.0%). As anticipated, with the exception of VEGF (2.0% T; 0.4% P) and svMP-i (1.7%; 12.6%) as part of the peptidic content, other lowly expressed ‘toxin’ families could not be assigned on the proteomic level.

The observed discrepancies in proteomic abundance and transcriptomic expression (e.g. CTLs and PLA_2_s) is influenced by many factors, e.g. post-genomic factors acting on toxin genes [49], such as the regulation of expression patterns by MicroRNAs (miRNA) [7,72], degradation processes [73], systematic or stochastic variations [74] or technical limitations in the experimental approach, including the eventually lower sensitivity of the proteomics workflow. Perhaps most importantly it needs to be mentioned that here we compared the toxin transcription level of a single individual (adult female) to a pooled venom protein sample (n=9), and thus, while it is possible that these differences are predominately due to the above mentioned regulatory processes, it seems likely that intra-specific venom variations may influence our findings. Due to understandable sampling/ethical restrictions relating to the sacrifice of individuals, we were unable to sequence venom gland transcriptomes of multiple specimens of *V. kaznakovi*.

The previous proteomic characterization of the *V. kaznakovi* venom by Utkin and coworkers was performed by in-solution trypsin proteolysis followed by nanoLC-MS/MS [52]. The PSM against a full NCBI Serpentes database identified 116 proteins from 14 typical snake venom protein families. The semi-quantitative venom composition showed PLA_2_ (41%) as the most abundant component, followed by svMPs (16%), CTL (12%), svSP (11%), CRISP (10%), LAAO (4%), VEGF (4%) and other lowly abundant proteins (< 1%) [52]. Besides the additional detection of lowly abundant proteins, the main difference to our results are the considerably higher levels of PLA2 and the lower abundance of svMPs (~ 4 fold difference for both protein families). The reasons for the additional detection of lowly abundant proteins could be of technical nature, as the nanoLC-MS/MS and mass spectrometer used in the study by Utkin *et al*., is typically more sensitive than the LC-MS/MS setup we used. While explanations for the major differences in protein abundance could be the different quantification method applied (UV abundance vs. summed peptide abundance [52]). Furthermore, the observed variations could also be biological in nature, i.e. the result of intra-specific venom variation, as the animals were collected in different geographic regions (Krasnodar Territory, Russia [51], with a distance of ~ 400 km to our collection site). However, as in most other venom proteomics studies the composition was determined from a pooled venom sample (15 individuals [52]), which has the potential to offset variation among individuals. In order to robustly assess the extent of intra-specific (e.g. population level) variations in *V. kaznakovi* venom analysis of a representative group of individuals is necessary.

### 3.4 Community venom profiling

It seems understandable that many venom studies are undertaken using pooled venom samples due to the associated costs and analysis time of decomplexing bottom-up venomics studies. In our case, we fractionated pooled venom from *V. kaznakovi* into 25 fractions and further separated the protein containing fractions (MW > 5 kDa) by SDS-PAGE. This multidimensional separation resulted in 25 digested peptide samples which we analyzed by LC-MS/MS, requiring ~ 10 h MS run time (25 min/sample), and an estimated ~$2,000 costs ($80/sample). Multiplying this effort and cost by numerous venom samples from individuals would of course make such a study comparatively expensive. Hence, many previous studies investigating venom variability within a species have used pooled venom for in-depth proteomic analysis, and then illuminated individual variability by the comparison of HPLC chromatograms and/or SDS-PAGE images [50,75,76]. This comparison allows at best a comparison at the protein family level (if protein families are clearly separated by HPLC or SDS-PAGE). As an alternative, a comparison by top-down or shotgun proteomics would allow for the differential comparison on the protein, or potentially proteoform, level performing a single LC-MS/MS run per individual.

However, shotgun approaches are likely to suffer from the aforementioned issues with protein inference, while top-down approaches have the drawback of not resolving high molecular mass proteins. This is particularly the case if the identification and comparison of proteins are based on Protein Spectrum Matching (PrSM), as high molecular weight toxins may not result in isotope resolved peaks and sufficient precursor signal, and thus are unlikely to provide sufficient fragment ions. However, a comparison by MS1 mass profiling only [77] would eliminate the problem of insufficient MS/MS fragments and isotope resolution, as spectra can be easily deconvoluted based on their charge state distribution. Such an approach could be particularly interesting for laboratories that are equipped with low resolution mass spectrometers.

In order to explore the potential of venom comparison by top-down mass profiling, we analyzed the venoms of nine *V. kaznakovi* individuals by LC-MS using the same chromatographic method as for our initial HPLC separation of our decomplexing bottom-up venom analysis. Chromatographic peak extraction of all individuals resulted in 119 consensus extracted ion chromatograms (XIC) or so-called ion features. The alignment of XICs by retention time and mass enabled the comparison of samples between individuals, but also a comparison with the mass profile of the pooled venom sample for a protein level annotation. An overview of all resulting features, including annotations, is shown in supplemental table 1. Looking at the binary distribution of ion features, individual venoms contained between 62 and 107 features, with a slightly higher average feature number in juveniles vs. adults. Comparing the total ion currents (TIC) of the LC-MS runs, the individual with the lowest feature number also had the lowest overall signal. Hence it is likely that the lower number of features in this individual was due to lower overall signal intensity and therefore might not be biologically representative. For further statistical evaluation we thus normalized feature abundance to TIC. Matching the features to the pooled bottom-up venomics results yielded an annotation rate between 83.4% and 93.5% of the features (based on XIC peak area). As anticipated, the annotation rate is slightly lower than the relative annotation of the pooled sample (96.0%; based on the UV_214_ peak area). The comparison of protein family venom compositions is shown in figure 4 and supplemental table 2. The highest variance was observed for svSP, CTL and LAAO toxin families (**Figure 5A**). Taking the age of the individuals into account, the abundance of svSPs was generally higher in the adult individuals than in the juveniles (average of 21.7% vs. 5.5 %), but no significant difference between male and female individuals, or between different geographic regions was observed. The svSPs play a significant role in mammalian envenomation by affecting the hemostatic system through perturbing blood coagulation, typically via the inducement of fibrinogenolytic effects [78,79]. Taking this into account, a possible explanation could be that lower svSP concentration in juveniles could be the result of differences in diet, as young animals typically prey on insects, before switching to feed upon small mammals and lizards as they become adults [80–82]. Despite their observed variations in abundance, no significant differences between the individual groups could be observed for the CTL and LAAO toxin families (**Figure 5A**). However, there was evidence that the svSP concentration is correlated to levels of LAAO, as the three individuals with the lowest svSP abundance showed the highest content of LAAO (**Figure 5A**). Whether this is a true biological effect or perhaps is the result of differences in ion suppression of the co-eluting compounds will need further investigation. We also observed variations between the PLA_2_ levels identified in the venoms, which ranged from 6.5–25.1%, but in all cases remained lower than those previously reported by Kovalchuk *et al*. (41%) [52]. In order to investigate the inner-species differences by multivariate statistics we performed a principal coordinate analysis (PCoA) using the Bray-Curtis dissimilarity metric. The PCoA plots of protein-level and proteoform-level data is shown in figure 5. Clustering of individuals in protein-family level PCoA space (**Figure 5B**) was only observed for the juvenile individuals. As expected from the univariate statistics no significant separation based on gender or region could be observed. Since an explanation for not resolving phenotype differences could be the reductions of variables through the binning of proteoforms, we used proteoform abundance as input matrix for PCoA. The outcomes of this analysis revealed a separation between both juvenile and adults, as well as between male and female snakes (**Figure 5C**). To investigating the toxin variants underpinning these separations, we used univariate comparison of the two groups and plotted the fold change of toxin abundance (log2) vs. the statistical significance (-log10 p-value, t-test) shown in supplemental figure 2. Besides the above mentioned differences in svSP, the most significant (p-value < 0.05, log2 fold change >2 or <-2) differences between juvenile and adult individuals was the higher abundance of small proteins with the masses 7707.26 Da, 5565.02 Da, 5693.10 Da in the juvenile group, all of which were unidentified in our proteomic analyses. Furthermore, we observed several smaller peptides with the masses 589.27 Da, 1244.56 Da, and 575.26 Da as well as a putative PLA_2_, with the mass of 13667.91 Da that was more abundant in the juveniles. Contrastingly, a putative PLA_2_ with a mass of 13683.86 Da was of lower abundance in the juvenile group. While we observed fewer significant changes between the venom toxins of the male and female individuals, the observed masses of the differential features indicated, that those differential toxins belong to different protein families than those involved in differentiating between juvenile and adult snakes. Two toxins with the masses 22829.66 Da and 24641.23 Da were higher abundant in male individuals and could be putatively annotated as hetero-dimeric CTLs. Another toxin with the mass 13549.87 was also higher abundant in the male group and according to the mass range is most likely a PLA_2_.

**Figure 4.**
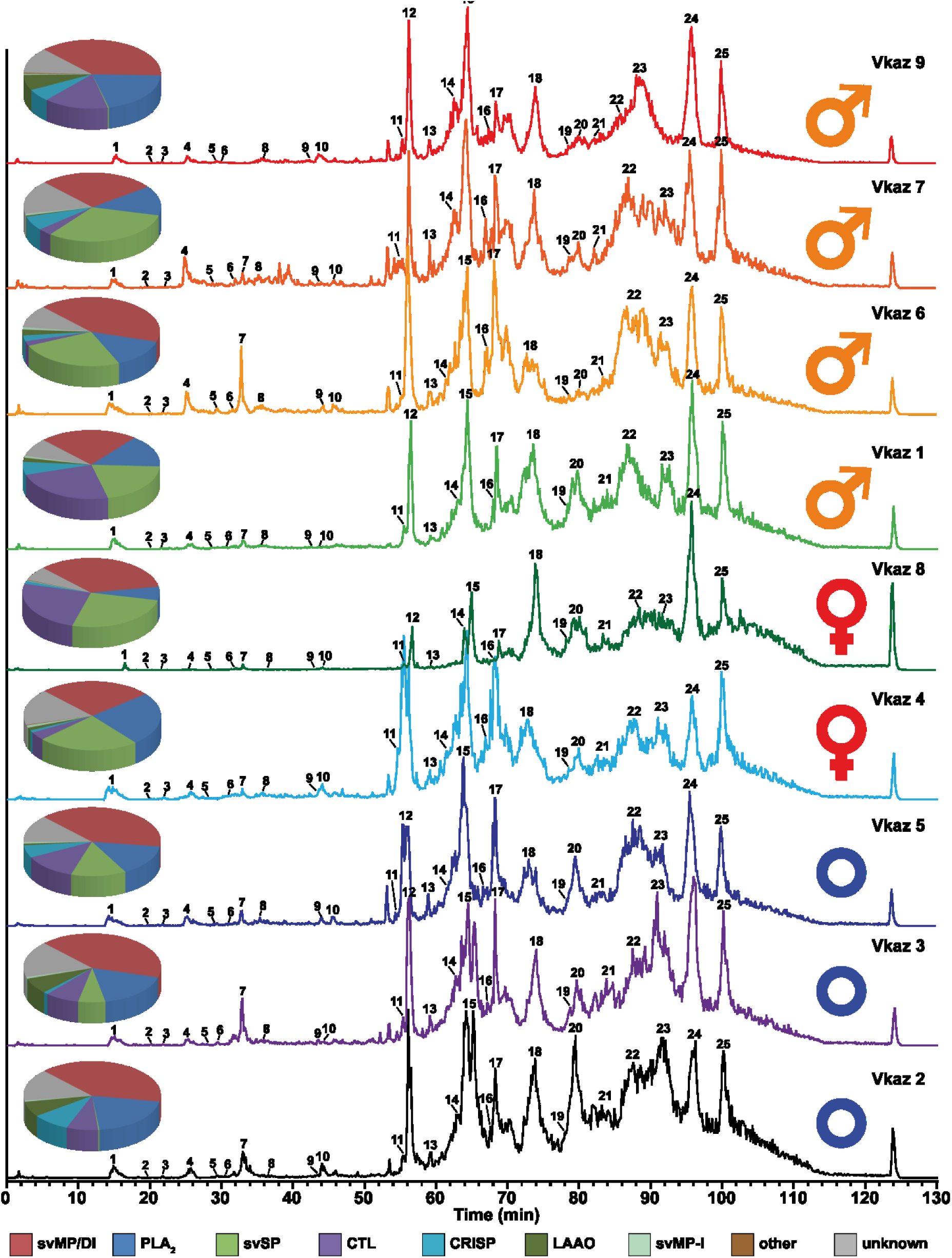
Intact molecular mass profiles of venom from several individuals of *V. kaznakovi*. The total ion counts (TIC) of native, crude venoms from several *V. kaznakovi* individuals were measured by HPLC-ESI-MS. The relative abundance was set to 100% for the highest peak. The peak nomenclature is based on the chromatogram fractions and is shown in figure 3A. The identified molecular masses of intact proteins and peptides are listed in supplemental table 2. The intact molecular mass profiling includes three juveniles of unknown sex (blue circle), and two female (red Venus symbol) and four male (orange Mars symbol) adult individuals.

**Figure 5.**
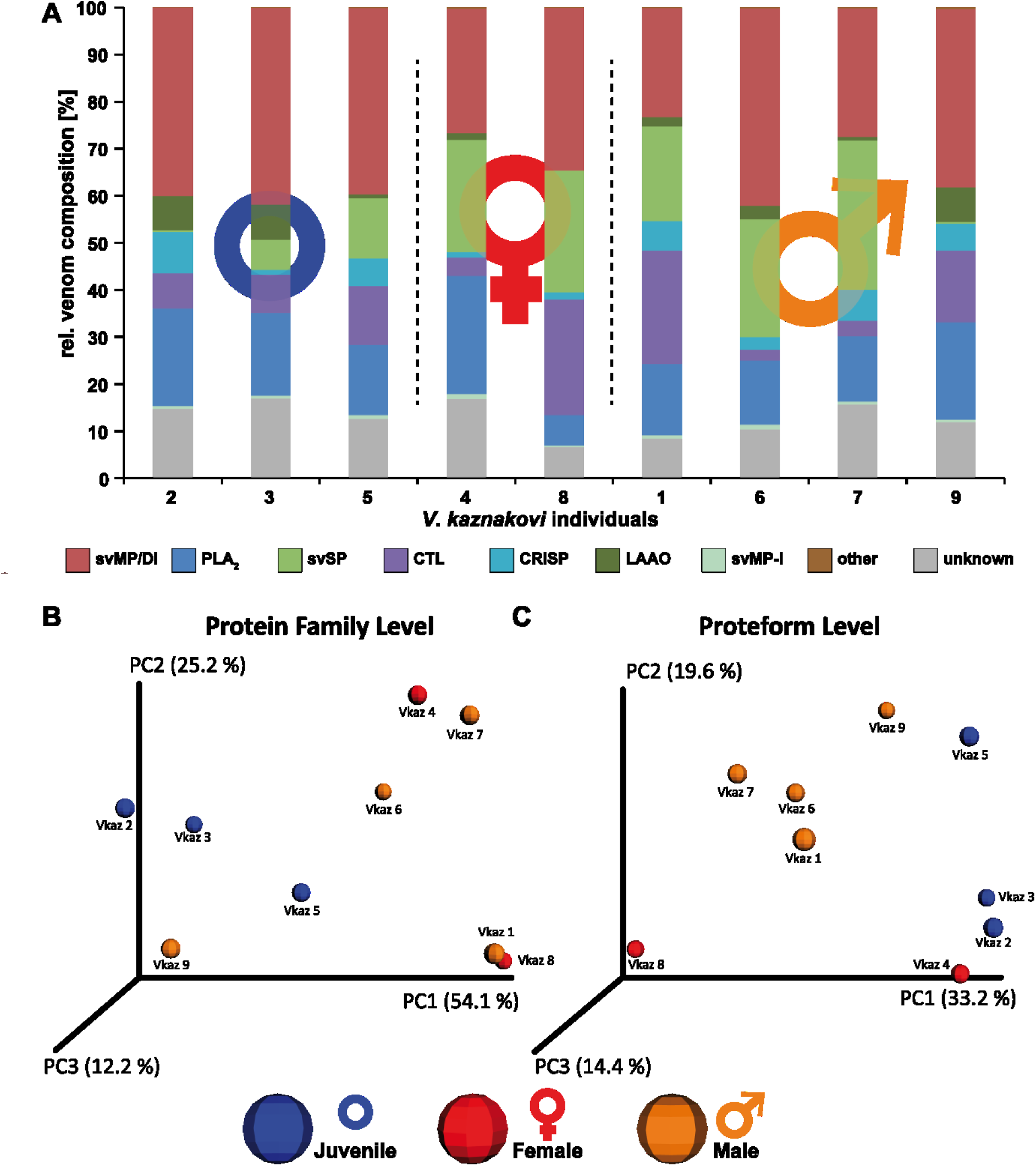
Principal Component Analysis (PCoA) and relative venom composition of individual *V. kaznakovi*. **A** The proteome overview includes three juveniles of unknown sex (blue circle), and two female (red Venus symbol) and four male (orange Mars symbol) adult individuals. The compositional similarity of venom is displayed through Bray-Curtis-Faith distance in PCoA space. Toxin similarity is visualized at the protein family level (**B**) and proteoform level (**C**).

## 4. Concluding remarks

Here we describe the detailed analysis of the venom composition of *Vipera kaznakovi* by a combination of venom gland transcriptomics and decomplexing bottom-up and top-down venom proteomics revealing the presence of 15 toxin families, of which the most abundant toxins were svMPs (37.7%), followed by PLA_2_s (19.0%), svSPs (9.5%), CTLs (6.9%) and CRISP (5.0%). Intact mass profiling enabled the rapid comparison of venom sourced from multiple individuals. This community venomics approach enabled higher sensitivity of direct intact protein analysis by LC-MS, in comparison to decomplexing bottom-up venomics, and thus enabled us working with multiple venom samples and with low amounts of material (< 0.5 mg venom). This allowed us to capture the snakes, perform venom extractions and then immediately release the animals back in to the field. Our approach revealed intraspecific venom variation in *Vipera kaznakovi*, including both ontogenetic differences between juvenile and adult snakes, and to a lesser extent, sexual differences between adult males and females. The highest significant difference in venom proteome composition was found between the adult and juvenile group, with svSP toxins found to exhibit the greatest variance. However, in addition, individuals within all groups showed a generally high relative variance of CTL and LAAO concentrations. svMPs on the other hand seemed to be constantly the most abundant venom component in all *V. kaznakovi* individuals analyzed in our study. However, as the statistical power with a relatively small subject size (n=9) is limited, it would be interesting to extend this study to a larger sample cohort, ideally covering all geographical regions (from Northeastern Turkey to Georgia and Russia) of the *V. kaznakovi* distribution zone. The workflow applied herein would be well suited for an extensive venom analysis at the population level, and will hopefully enable venom researchers to more easily expand their experimental approach towards robust comparisons of intra-species venom variation, and not only characterize pooled venom samples.

## Author contributions

D.P., A.N. and R.D.S. planned the study. D.P., A.N., B.G., M.K. and P.H. collected the animals and prepared venom and venom gland tissue samples. A.N. performed the determination of acute lethal dose. D.P., P.H. and B.-F.H. performed the toxin separation and acquired the mass spectrometry data. G.W., S.C.W. and N.R.C. constructed the transcriptome. D.P., B.-F.H. and N.R.C. performed the data analysis. A.N., N.R.C. and R.D.S. acquired funding and provided materials and instruments for the study. D.P., B.-F.H. and R.D.S. wrote the manuscript. All authors read, discussed and approved the manuscript.

## Acknowledgements

We thank Sabah Ul-Hasan and Anthony Saviola for critical reading of the paper draft, and Robert Harrison for assistance with transcriptomics. This study was supported by the Deutsche Forschungsgemeinschaft (DFG) through the Cluster of Excellence ‘Unifying Concepts in Catalysis (UniCat), a postdoctoral research scholarship to D.P. (PE 2600/1), the Scientific and Technical Research Council of Turkey (TÜBİTAK) under Grant 114Z946, and a Sir Henry Dale Fellowship (200517/Z/16/Z) jointly funded by the Wellcome Trust and the Royal Society to N.R.C

---

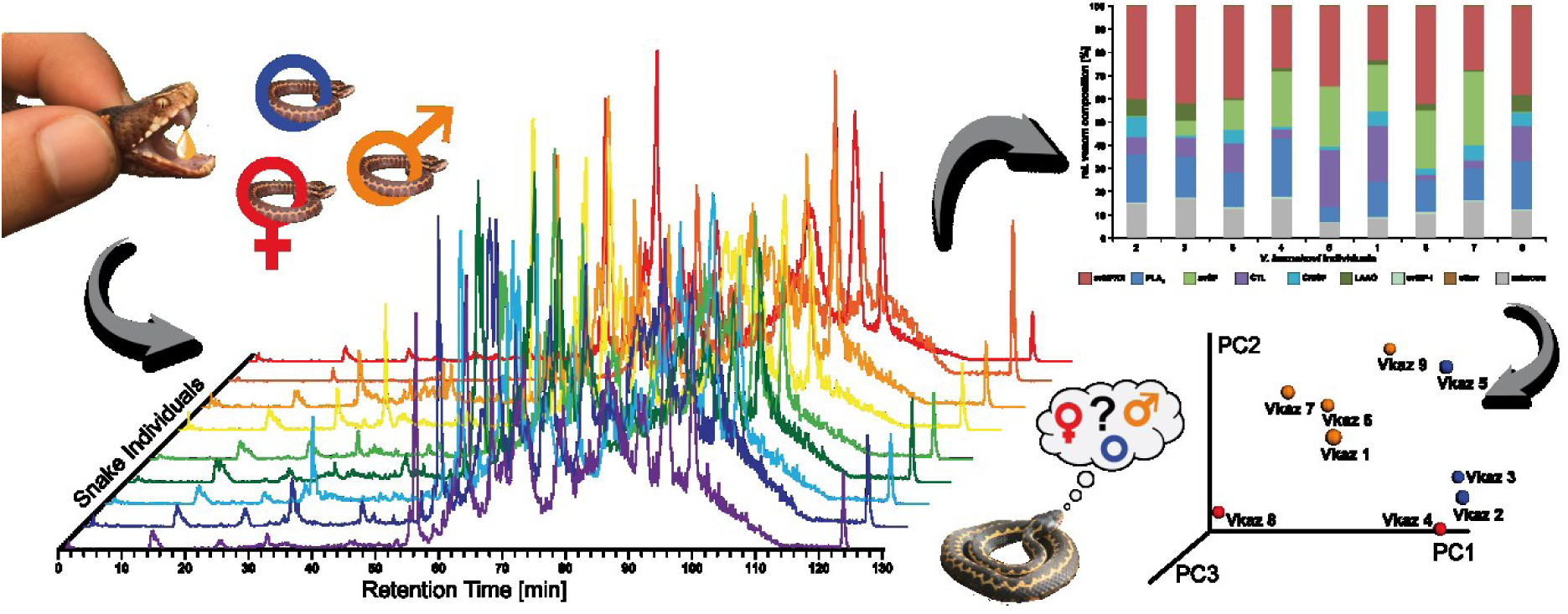
Graphical Abstract.

## References

[1] Calvete JJ, Lomonte B. A bright future for integrative venomics. Toxicon 2015;107(Pt B):159–62.

[2] Juárez P, Sanz L, Calvete JJ. Snake venomics: Characterization of protein families in Sistrurus barbouri venom by cysteine mapping, N-terminal sequencing, and tandem mass spectrometry analysis. Proteomics 2004;4(2):327–38.

[3] Calvete JJ. Snake venomics - from low-resolution toxin-pattern recognition to toxin-resolved venom proteomes with absolute quantification. Expert Rev Proteomics 2018;15(7):555–68.

[4] Vonk FJ, Casewell NR, Henkel CV, Heimberg AM, Jansen HJ, McCleary RJR et al. The king cobra genome reveals dynamic gene evolution and adaptation in the snake venom system. Proc Natl Acad Sci U S A 2013;110(51):20651–6.

[5] Aird SD, Arora J, Barua A, Qiu L, Terada K, Mikheyev AS. Population Genomic Analysis of a Pitviper Reveals Microevolutionary Forces Underlying Venom Chemistry. Genome Biol Evol 2017;9(10):2640–9.

[6] Pla D, Petras D, Saviola AJ, Modahl CM, Sanz L, Pérez A et al. Transcriptomics-guided bottom-up and top-down venomics of neonate and adult specimens of the arboreal rear-fanged Brown Treesnake, Boiga irregularis, from Guam. J Proteomics 2018;174:71–84.

[7] Durban J, Sanz L, Trevisan-Silva D, Neri-Castro E, Alagón A, Calvete JJ. Integrated Venomics and Venom Gland Transcriptome Analysis of Juvenile and Adult Mexican Rattlesnakes Crotalus simus, C. tzabcan, and C. culminatus Revealed miRNA-modulated Ontogenetic Shifts. J Proteome Res 2017;16(9):3370–90.

[8] Aird SD, da Silva NJ, Qiu L, Villar-Briones A, Saddi VA, Pires de Campos Telles M et al. Coralsnake Venomics: Analyses of Venom Gland Transcriptomes and Proteomes of Six Brazilian Taxa. Toxins (Basel) 2017;9(6).

[9] Pla D, Sanz L, Whiteley G, Wagstaff SC, Harrison RA, Casewell NR et al. What killed Karl Patterson Schmidt? Combined venom gland transcriptomic, venomic and antivenomic analysis of the South African green tree snake (the boomslang), Dispholidus typus. Biochim Biophys Acta 2017;1861(4):814–23.

[10] Gonçalves-Machado L, Pla D, Sanz L, Jorge RJB, Leitão-De-Araújo M, Alves MLM et al. Combined venomics, venom gland transcriptomics, bioactivities, and antivenomics of two Bothrops jararaca populations from geographic isolated regions within the Brazilian Atlantic rainforest. J Proteomics 2016;135:73–89.

[11] Fry BG, Scheib H, L M Junqueira Azevedo I de, Silva DA, Casewell NR. Novel transcripts in the maxillary venom glands of advanced snakes. Toxicon 2012;59(7–8):696–708.

[12] Casewell NR, Harrison RA, Wüster W, Wagstaff SC. Comparative venom gland transcriptome surveys of the saw-scaled vipers (Viperidae: Echis) reveal substantial intra-family gene diversity and novel venom transcripts. BMC Genomics 2009;10:564.

[13] Rokyta DR, Wray KP, Margres MJ. The genesis of an exceptionally lethal venom in the timber rattlesnake (Crotalus horridus) revealed through comparative venom-gland transcriptomics. BMC Genomics 2013;14:394.

[14] Tan CH, Tan KY, Fung SY, Tan NH. Venom-gland transcriptome and venom proteome of the Malaysian king cobra (Ophiophagus hannah). BMC Genomics 2015;16:687.

[15] Nawarak J, Sinchaikul S, Wu C-Y, Liau M-Y, Phutrakul S, Chen S-T. Proteomics of snake venoms from Elapidae and Viperidae families by multidimensional chromatographic methods. Electrophoresis 2003;24(16):2838–54.

[16] Tasoulis T, Isbister GK. A Review and Database of Snake Venom Proteomes. Toxins (Basel) 2017;9(9).

[17] Lomonte B, Calvete JJ. Strategies in ‘snake venomics’ aiming at an integrative view of compositional, functional, and immunological characteristics of venoms. J Venom Anim Toxins Incl Trop Dis 2017;23:26.

[18] Melani RD, Goto-Silva L, Nogueira FCS, Junqueira M, Domont GB. Shotgun Approaches for Venom Analysis. In: Gopalakrishnakone P, Calvete JJ, editors. Venom Genomics and Proteomics: Venom Genomics and Proteomics. Dordrecht: Springer Netherlands; 2014, p. 1–12.

[19] Nesvizhskii AI, Aebersold R. Interpretation of shotgun proteomic data: The protein inference problem. Mol Cell Proteomics 2005;4(10):1419–40.

[20] Smith LM, Kelleher NL. Proteoform: A single term describing protein complexity. Nat Methods 2013;10(3):186–7.

[21] Petras D, Heiss P, Süssmuth RD, Calvete JJ. Venom Proteomics of Indonesian King Cobra, Ophiophagus hannah: Integrating Top-Down and Bottom-Up Approaches. J Proteome Res 2015;14(6):2539–56.

[22] Melani RD, Skinner OS, Fornelli L, Domont GB, Compton PD, Kelleher NL. Mapping Proteoforms and Protein Complexes From King Cobra Venom Using Both Denaturing and Native Top-down Proteomics. Mol Cell Proteomics 2016;15(7):2423–34.

[23] Petras D, Heiss P, Harrison RA, Süssmuth RD, Calvete JJ. Top-down venomics of the East African green mamba, Dendroaspis angusticeps, and the black mamba, Dendroaspis polylepis, highlight the complexity of their toxin arsenals. J Proteomics 2016;146:148–64.

[24] Ainsworth S, Petras D, Engmark M, Süssmuth RD, Whiteley G, Albulescu L-O et al. The medical threat of mamba envenoming in sub-Saharan Africa revealed by genus-wide analysis of venom composition, toxicity and antivenomics profiling of available antivenoms. J Proteomics 2018;172:173–89.

[25] Damm M, Hempel B-F, Nalbantsoy A, Süssmuth RD. Comprehensive Snake Venomics of the Okinawa Habu Pit Viper, Protobothrops flavoviridis, by Complementary Mass Spectrometry-Guided Approaches. Molecules 2018;23(8).

[26] Göçmen B, Heiss P, Petras D, Nalbantsoy A, Süssmuth RD. Mass spectrometry guided venom profiling and bioactivity screening of the Anatolian Meadow Viper, Vipera anatolica. Toxicon 2015;107(Pt B):163–74.

[27] Hempel B-F, Damm M, Göçmen B, Karis M, Oguz MA, Nalbantsoy A et al. Comparative Venomics of the Vipera ammodytes transcaucasiana and Vipera ammodytes montandoni from Turkey Provides Insights into Kinship. Toxins (Basel) 2018;10(1).

[28] Nalbantsoy A, Hempel B-F, Petras D, Heiss P, Göçmen B, Iğci N et al. Combined venom profiling and cytotoxicity screening of the Radde’s mountain viper (Montivipera raddei) and Mount Bulgar Viper (Montivipera bulgardaghica) with potent cytotoxicity against human A549 lung carcinoma cells. Toxicon 2017;135:71–83.

[29] Ghezellou P, Garikapati V, Kazemi SM, Strupat K, Ghassempour A, Spengler B. A perspective view of top-down proteomics in snake venom research. Rapid Commun Mass Spectrom 2018.

[30] Wu S, Brown JN, Tolić N, Meng D, Liu X, Zhang H et al. Quantitative analysis of human salivary gland-derived intact proteome using top-down mass spectrometry. Proteomics 2014;14(10):1211–22.

[31] Moehring F, Waas M, Keppel TR, Rathore D, Cowie AM, Stucky CL et al. Quantitative Top-Down Mass Spectrometry Identifies Proteoforms Differentially Released during Mechanical Stimulation of Mouse Skin. J Proteome Res 2018;17(8):2635–48.

[32] Ntai I, Toby TK, LeDuc RD, Kelleher NL. A Method for Label-Free, Differential Top-Down Proteomics. Methods Mol Biol 2016;1410:121–33.

[33] Daltry JC, Wüster W, Thorpe RS. Diet and snake venom evolution. Nature 1996;379(6565):537–40.

[34] Barlow A, Pook CE, Harrison RA, Wüster W. Coevolution of diet and prey-specific venom activity supports the role of selection in snake venom evolution. Proc Biol Sci 2009;276(1666):2443–9.

[35] Gibbs HL, Mackessy SP. Functional basis of a molecular adaptation: Prey-specific toxic effects of venom from Sistrurus rattlesnakes. Toxicon 2009;53(6):672–9.

[36] Gibbs HL, Sanz L, Sovic MG, Calvete JJ. Phylogeny-based comparative analysis of venom proteome variation in a clade of rattlesnakes (Sistrurus sp.). PLoS ONE 2013;8(6):e67220.

[37] Alape-Girón A, Sanz L, Escolano J, Flores-Díaz M, Madrigal M, Sasa M et al. Snake venomics of the lancehead pitviper Bothrops asper: Geographic, individual, and ontogenetic variations. J Proteome Res 2008;7(8):3556–71.

[38] Huang H-W, Liu B-S, Chien K-Y, Chiang L-C, Huang S-Y, Sung W-C et al. Cobra venom proteome and glycome determined from individual snakes of Naja atra reveal medically important dynamic range and systematic geographic variation. J Proteomics 2015;128:92–104.

[39] Jorge RJB, Monteiro HSA, Gonçalves-Machado L, Guarnieri MC, Ximenes RM, Borges-Nojosa DM et al. Venomics and antivenomics of Bothrops erythromelas from five geographic populations within the Caatinga ecoregion of northeastern Brazil. J Proteomics 2015;114:93–114.

[40] Pimenta DC, Prezoto BC, Konno K, Melo RL, Furtado MF, Camargo ACM et al. Mass spectrometric analysis of the individual variability of Bothrops jararaca venom peptide fraction. Evidence for sex-based variation among the bradykinin-potentiating peptides. Rapid Commun Mass Spectrom 2007;21(6):1034–42.

[41] Menezes MC, Furtado MF, Travaglia-Cardoso SR, Camargo ACM, Serrano SMT. Sex-based individual variation of snake venom proteome among eighteen Bothrops jararaca siblings. Toxicon 2006;47(3):304–12.

[42] Amorim FG, Costa TR, Baiwir D, Pauw E de, Quinton L, Sampaio SV. Proteopeptidomic, Functional and Immunoreactivity Characterization of Bothrops moojeni Snake Venom: Influence of Snake Gender on Venom Composition. Toxins (Basel) 2018;10(5).

[43] Mackessy SP. Venom Ontogeny in the Pacific Rattlesnakes Crotalus viridis helleri and C. v. oreganus. Copeia 1988;1988(1):92.

[44] Gutiérrez JM, Avila C, Camacho Z, Lomonte B. Ontogenetic changes in the venom of the snake Lachesis muta stenophrys (bushmaster) from Costa Rica. Toxicon 1990;28(4):419–26.

[45] Guércio RAP, Shevchenko A, Shevchenko A, López-Lozano JL, Paba J, Sousa MV et al. Ontogenetic variations in the venom proteome of the Amazonian snake Bothrops atrox. Proteome Sci 2006;4:11.

[46] Mackessy SP, Sixberry NM, Heyborne WH, Fritts T. Venom of the Brown Treesnake, Boiga irregularis: Ontogenetic shifts and taxa-specific toxicity. Toxicon 2006;47(5):537–48.

[47] Calvete JJ, Casewell NR, Hernández-Guzmán U, Quesada-Bernat S, Sanz L, Rokyta DR et al. Venom Complexity in a Pitviper Produced by Facultative Parthenogenesis. Sci Rep 2018;8(1):11539.

[48] Casewell NR, Wüster W, Vonk FJ, Harrison RA, Fry BG. Complex cocktails: The evolutionary novelty of venoms. Trends Ecol Evol (Amst) 2013;28(4):219–29.

[49] Casewell NR, Wagstaff SC, Wüster W, Cook DAN, Bolton FMS, King SI et al. Medically important differences in snake venom composition are dictated by distinct postgenomic mechanisms. Proc Natl Acad Sci U S A 2014;111(25):9205–10.

[50] Massey DJ, Calvete JJ, Sánchez EE, Sanz L, Richards K, Curtis R et al. Venom variability and envenoming severity outcomes of the Crotalus scutulatus scutulatus (Mojave rattlesnake) from Southern Arizona. J Proteomics 2012;75(9):2576–87.

[51] Starkov VG, Osipov AV, Utkin YN. Toxicity of venoms from vipers of Pelias group to crickets Gryllus assimilis and its relation to snake entomophagy. Toxicon 2007;49(7):995–1001.

[52] Kovalchuk SI, Ziganshin RH, Starkov VG, Tsetlin VI, Utkin YN. Quantitative Proteomic Analysis of Venoms from Russian Vipers of Pelias Group: Phospholipases A_2_ are the Main Venom Components. Toxins (Basel) 2016;8(4):105.

[53] Bruce RD. An up-and-down procedure for acute toxicity testing. Fundam Appl Toxicol 1985;5(1):151–7.

[54] Cates CC, McCabe JG, Lawson GW, Couto MA. Core body temperature as adjunct to endpoint determination in murine median lethal dose testing of rattlesnake venom. Comp Med 2014;64(6):440–7.

[55] Ethical principles and guidelines for experiments on animals. Swiss Academy of Medical Sciences. Swiss Academy of Sciences. Experientia 1996;52(1):1–3.

[56] Archer J, Whiteley G, Casewell NR, Harrison RA, Wagstaff SC. VTBuilder: A tool for the assembly of multi isoform transcriptomes. BMC Bioinformatics 2014;15:389.

[57] Götz S, García-Gómez JM, Terol J, Williams TD, Nagaraj SH, Nueda MJ et al. High-throughput functional annotation and data mining with the Blast2GO suite. Nucleic Acids Res 2008;36(10):3420–35.

[58] Muth T, Weilnböck L, Rapp E, Huber CG, Martens L, Vaudel M et al. DeNovoGUI: An open source graphical user interface for de novo sequencing of tandem mass spectra. J Proteome Res 2014;13(2):1143–6.

[59] Altschul SF, Gish W, Miller W, Myers EW, Lipman DJ. Basic local alignment search tool. J Mol Biol 1990;215(3):403–10.

[60] Barsnes H, Vaudel M. SearchGUI: A Highly Adaptable Common Interface for Proteomics Search and de Novo Engines. J Proteome Res 2018;17(7):2552–5.

[61] Vizcaíno JA, Deutsch EW, Wang R, Csordas A, Reisinger F, Ríos D et al. ProteomeXchange provides globally coordinated proteomics data submission and dissemination. Nat Biotechnol 2014;32(3):223–6.

[62] Sket D, Gubensek F, Adamic S, Lebez D. Action of a partially purified basic protein fraction from Vipera ammodytes venom. Toxicon 1973;11(1):47–53.

[63] Oukkache N, El Jaoudi R, Ghalim N, Chgoury F, Bouhaouala B, Mdaghri NE et al. Evaluation of the lethal potency of scorpion and snake venoms and comparison between intraperitoneal and intravenous injection routes. Toxins (Basel) 2014;6(6):1873–81.

[64] Calderón L, Lomonte B, Gutiérrez JM, Tarkowski A, Hanson LA. Biological and biochemical activities of Vipera berus (European viper) venom. Toxicon 1993;31(6):743–53.

[65] Haro L de, Robbe-Vincent A, Saliou B, Valli M, Bon C, Choumet V. Unusual neurotoxic envenomations by Vipera aspis aspis snakes in France. Hum Exp Toxicol 2002;21(3):137–45.

[66] Rokyta DR, Lemmon AR, Margres MJ, Aronow K. The venom-gland transcriptome of the eastern diamondback rattlesnake (Crotalus adamanteus). BMC Genomics 2012;13:312.

[67] Junqueira-de-Azevedo ILM, Bastos CMV, Ho PL, Luna MS, Yamanouye N, Casewell NR. Venom-related transcripts from Bothrops jararaca tissues provide novel molecular insights into the production and evolution of snake venom. Mol Biol Evol 2015;32(3):754–66.

[68] Fox JW, Serrano SMT. Insights into and speculations about snake venom metalloproteinase (SVMP) synthesis, folding and disulfide bond formation and their contribution to venom complexity. FEBS J 2008;275(12):3016–30.

[69] Casewell NR, Wagstaff SC, Harrison RA, Renjifo C, Wüster W. Domain loss facilitates accelerated evolution and neofunctionalization of duplicate snake venom metalloproteinase toxin genes. Mol Biol Evol 2011;28(9):2637–49.

[70] Gasmi A, Srairi N, Guermazi S, Dekhil H, Dkhil H, Karoui H et al. Amino acid structure and characterization of a heterodimeric disintegrin from Vipera lebetina venom. Biochim Biophys Acta 2001;1547(1):51–6.

[71] Navdaev A, Clemetson JM, Polgar J, Kehrel BE, Glauner M, Magnenat E et al. Aggretin, a heterodimeric C-type lectin from Calloselasma rhodostoma (Malayan pit viper), stimulates platelets by binding to α2β1 integrin and glycoprotein Ib, activating Syk and phospholipase Cγ 2, but does not involve the glycoprotein VI/Fc receptor γ chain collagen receptor. J Biol Chem 2001;276(24):20882–9.

[72] Durban J, Pérez A, Sanz L, Gómez A, Bonilla F, Rodríguez S et al. Integrated “omics” profiling indicates that miRNAs are modulators of the ontogenetic venom composition shift in the Central American rattlesnake, Crotalus simus simus. BMC Genomics 2013;14:234.

[73] Vogel C, Marcotte EM. Insights into the regulation of protein abundance from proteomic and transcriptomic analyses. Nat Rev Genet 2012;13(4):227–32.

[74] Ruggles KV, Krug K, Wang X, Clauser KR, Wang J, Payne SH et al. Methods, Tools and Current Perspectives in Proteogenomics. Mol Cell Proteomics 2017;16(6):959–81.

[75] Calvete JJ, Sanz L, Pérez A, Borges A, Vargas AM, Lomonte B et al. Snake population venomics and antivenomics of Bothrops atrox: Paedomorphism along its transamazonian dispersal and implications of geographic venom variability on snakebite management. J Proteomics 2011;74(4):510–27.

[76] Galizio NdC, Serino-Silva C, Stuginski DR, Abreu PAE, Sant’Anna SS, Grego KF et al. Compositional and functional investigation of individual and pooled venoms from long-term captive and recently wild-caught Bothrops jararaca snakes. J Proteomics 2018;186:56–70.

[77] Fry BG, Wickramaratna JC, Hodgson WC, Alewood PF, Kini RM, Ho H et al. Electrospray liquid chromatography/mass spectrometry fingerprinting of Acanthophis (death adder) venoms: Taxonomic and toxinological implications. Rapid Commun Mass Spectrom 2002;16(6):600–8.

[78] Serrano SMT. The long road of research on snake venom serine proteinases. Toxicon 2013;62:19–26.

[79] Slagboom J, Kool J, Harrison RA, Casewell NR. Haemotoxic snake venoms: Their functional activity, impact on snakebite victims and pharmaceutical promise. Br J Haematol 2017;177(6):947–59.

[80] Dias GS, Kitano ES, Pagotto AH, Sant’anna SS, Rocha MMT, Zelanis A et al. Individual variability in the venom proteome of juvenile Bothrops jararaca specimens. J Proteome Res 2013;12(10):4585–98.

[81] Gibbs HL, Sanz L, Chiucchi JE, Farrell TM, Calvete JJ. Proteomic analysis of ontogenetic and diet-related changes in venom composition of juvenile and adult Dusky Pigmy rattlesnakes (Sistrurus miliarius barbouri). J Proteomics 2011;74(10):2169–79.

[82] Zelanis A, Tashima AK, Rocha MMT, Furtado MF, Camargo ACM, Ho PL et al. Analysis of the ontogenetic variation in the venom proteome/peptidome of Bothrops jararaca reveals different strategies to deal with prey. J Proteome Res 2010;9(5):2278–91.

[83] Geniez P. Snakes of Europe, North Africa & the Middle East: A photographic guide. Princeton: Princeton University Press; 2018.

