## Supplemental Information for "Community venomics reveals intra-species variations in venom composition of a local population of *Vipera kaznakovi* in Northeastern Turkey"

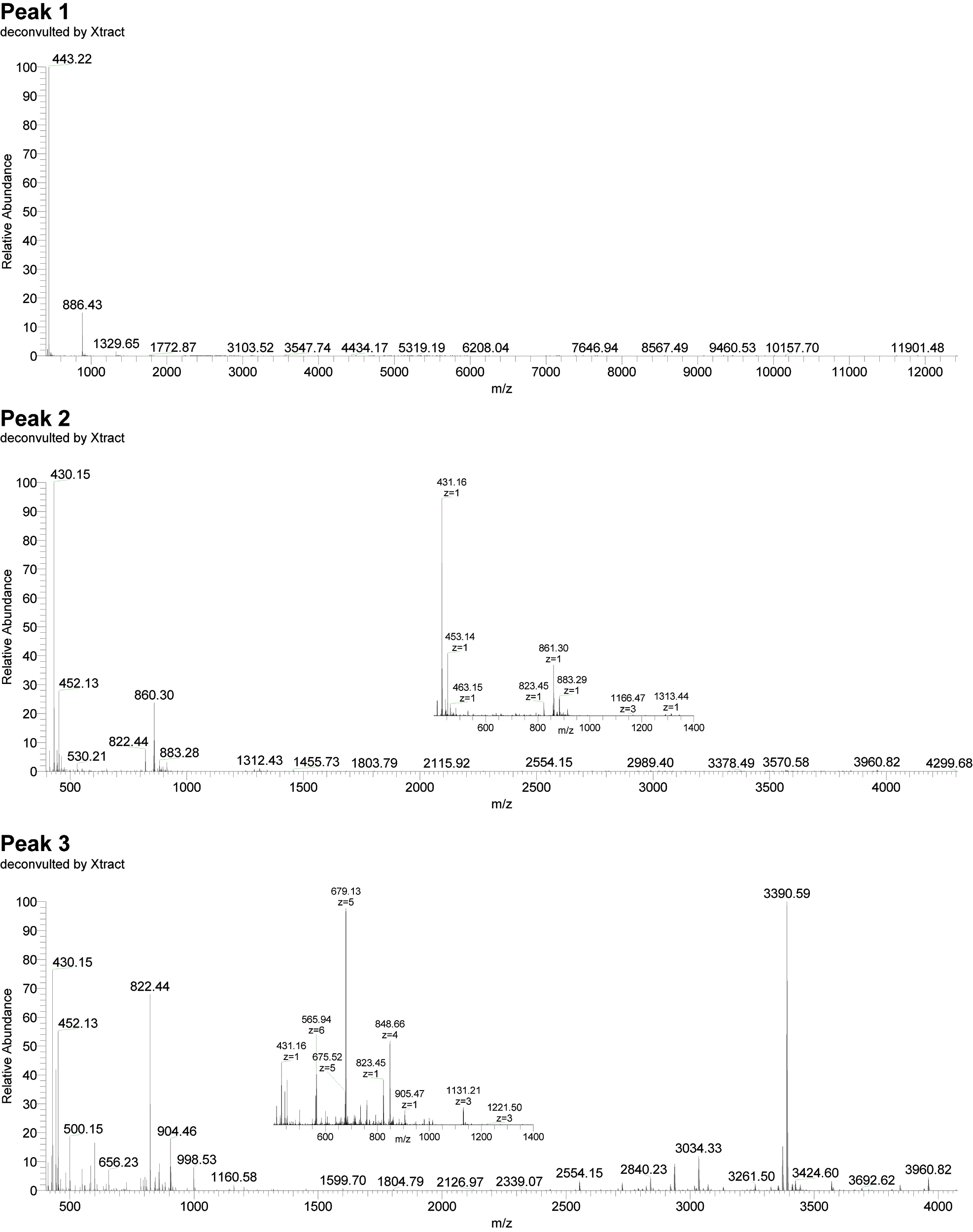

**Supplemental Figure 1. Intact mass spectra of pooled Vipera kaznakovi venom**. Peak nomenclature is based on the chromatogram fractions (see **Figure 3**). Mass spectra were either isotopically deconvoluted with Xcalibur or charge deconvoluted with magic transformer (MagTran).

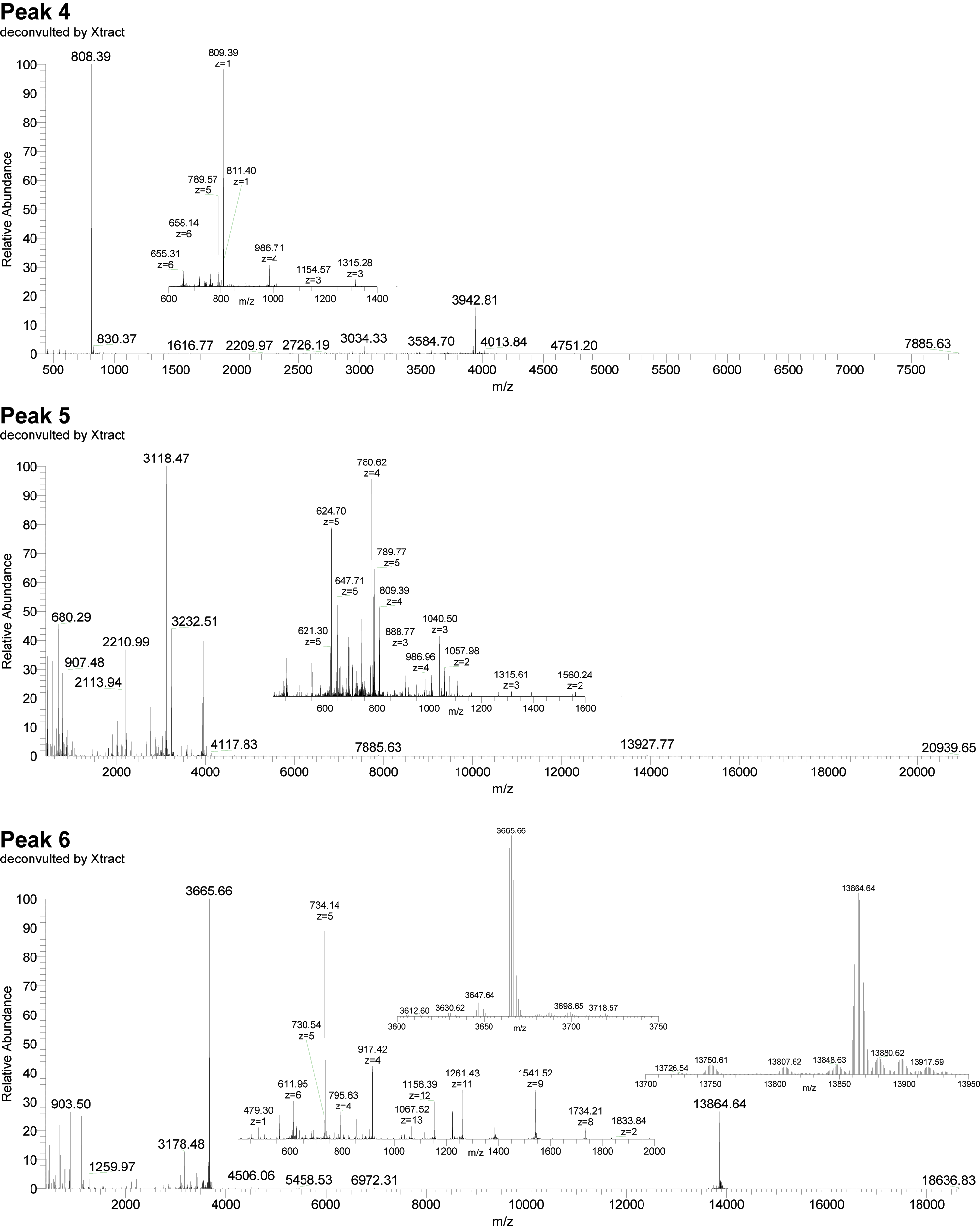

**Supplemental Figure 1. continued**

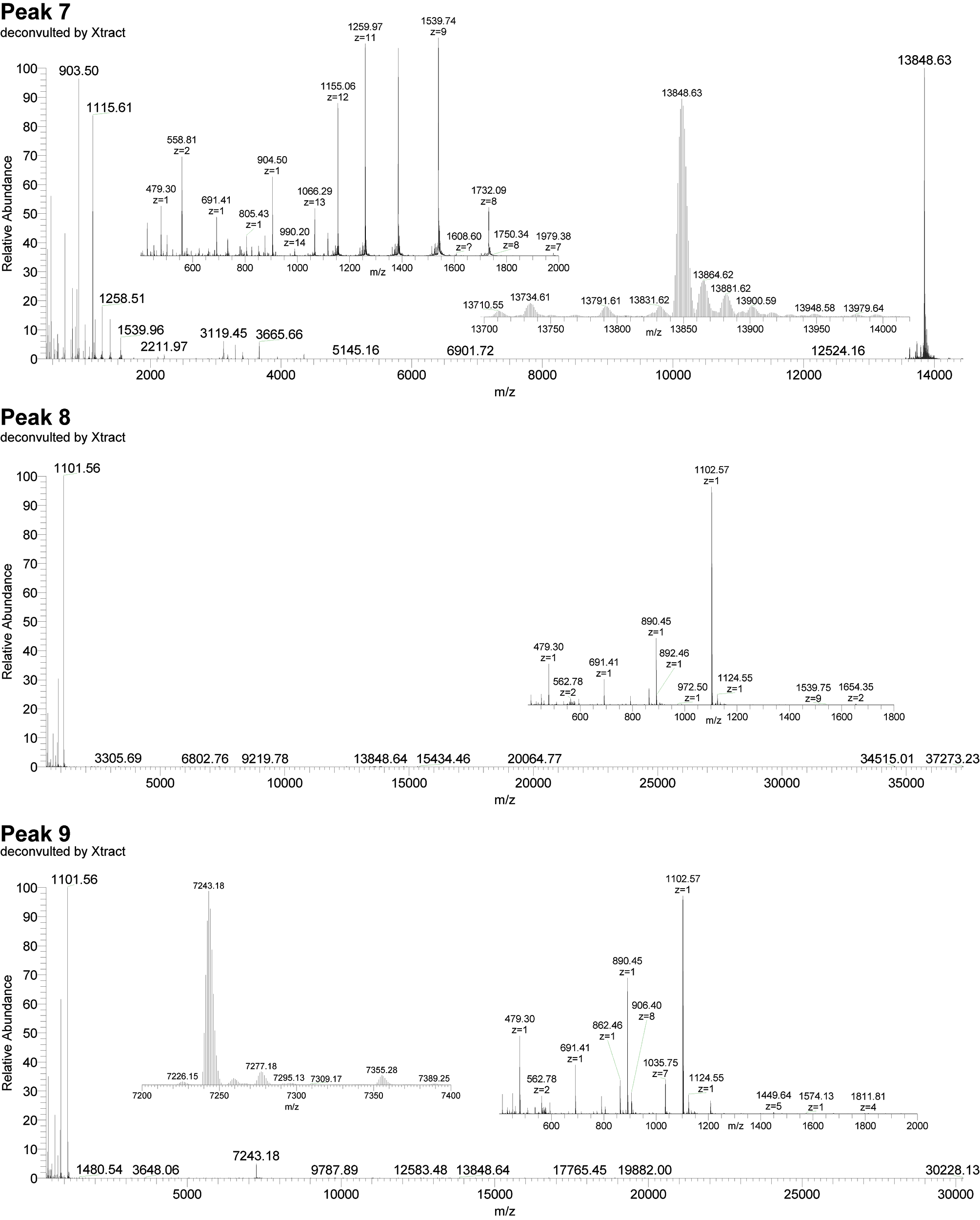

**Supplemental Figure 1. continued**

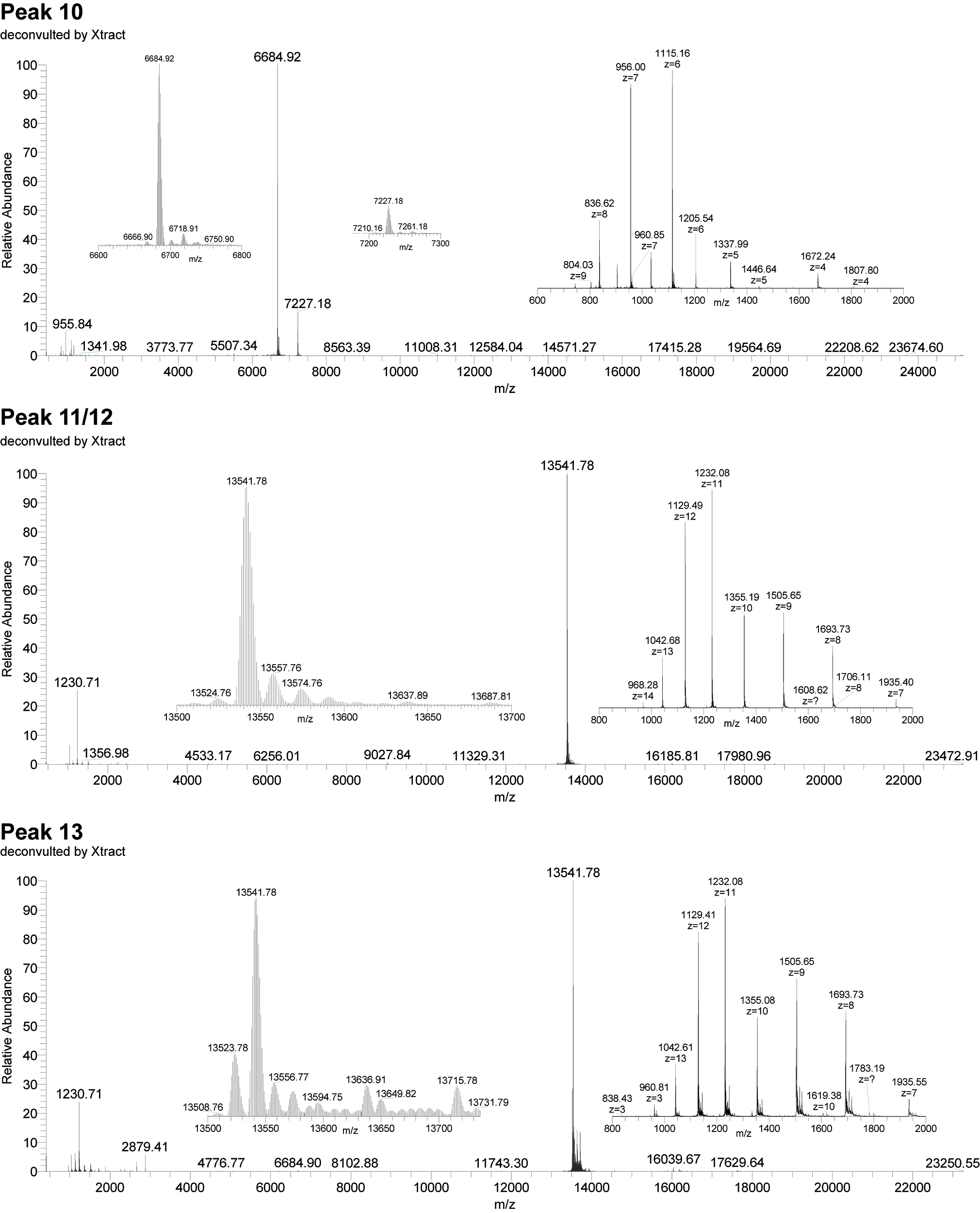

**Supplemental Figure 1. continued**

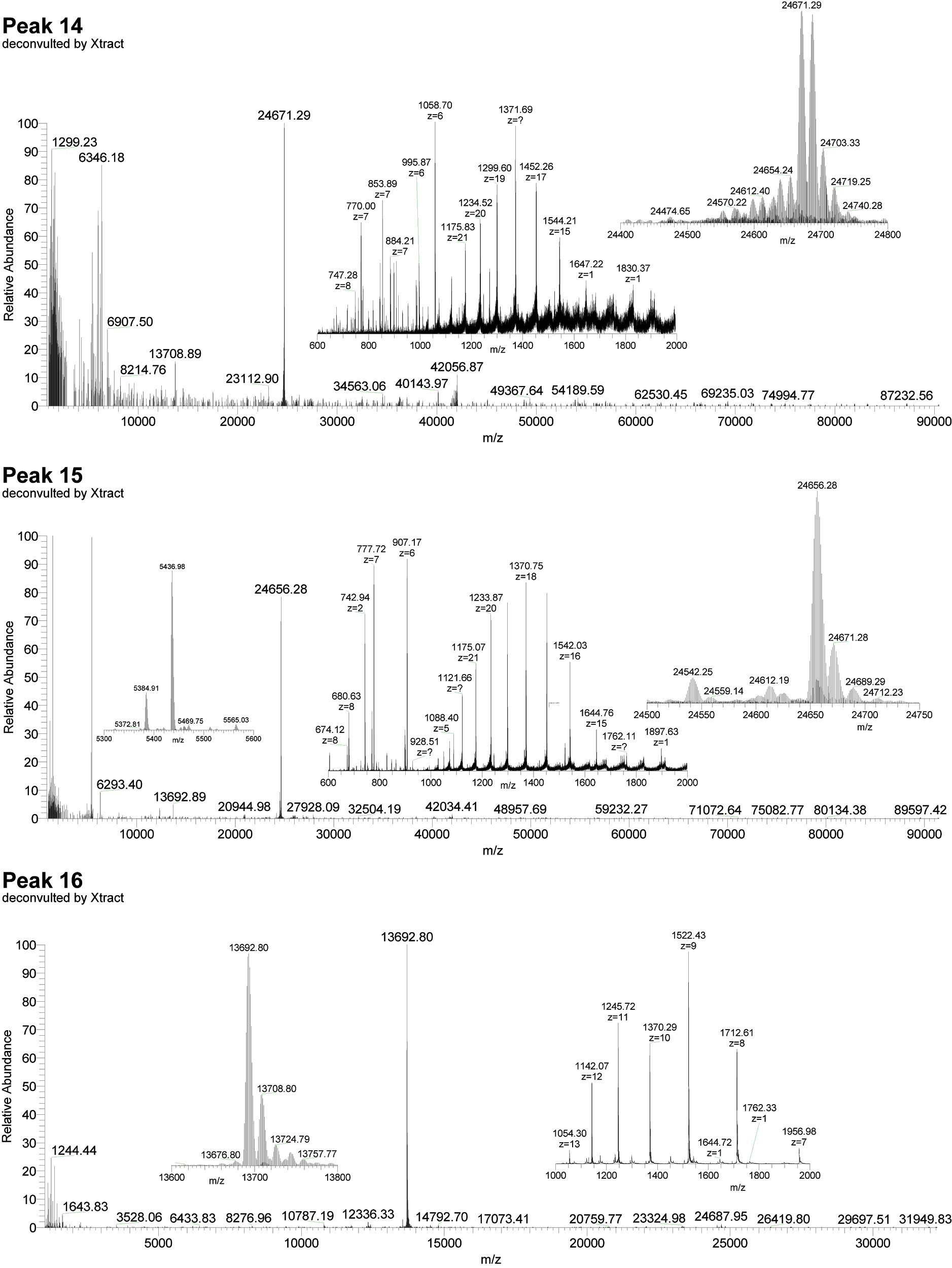

**Supplemental Figure 1. continued**

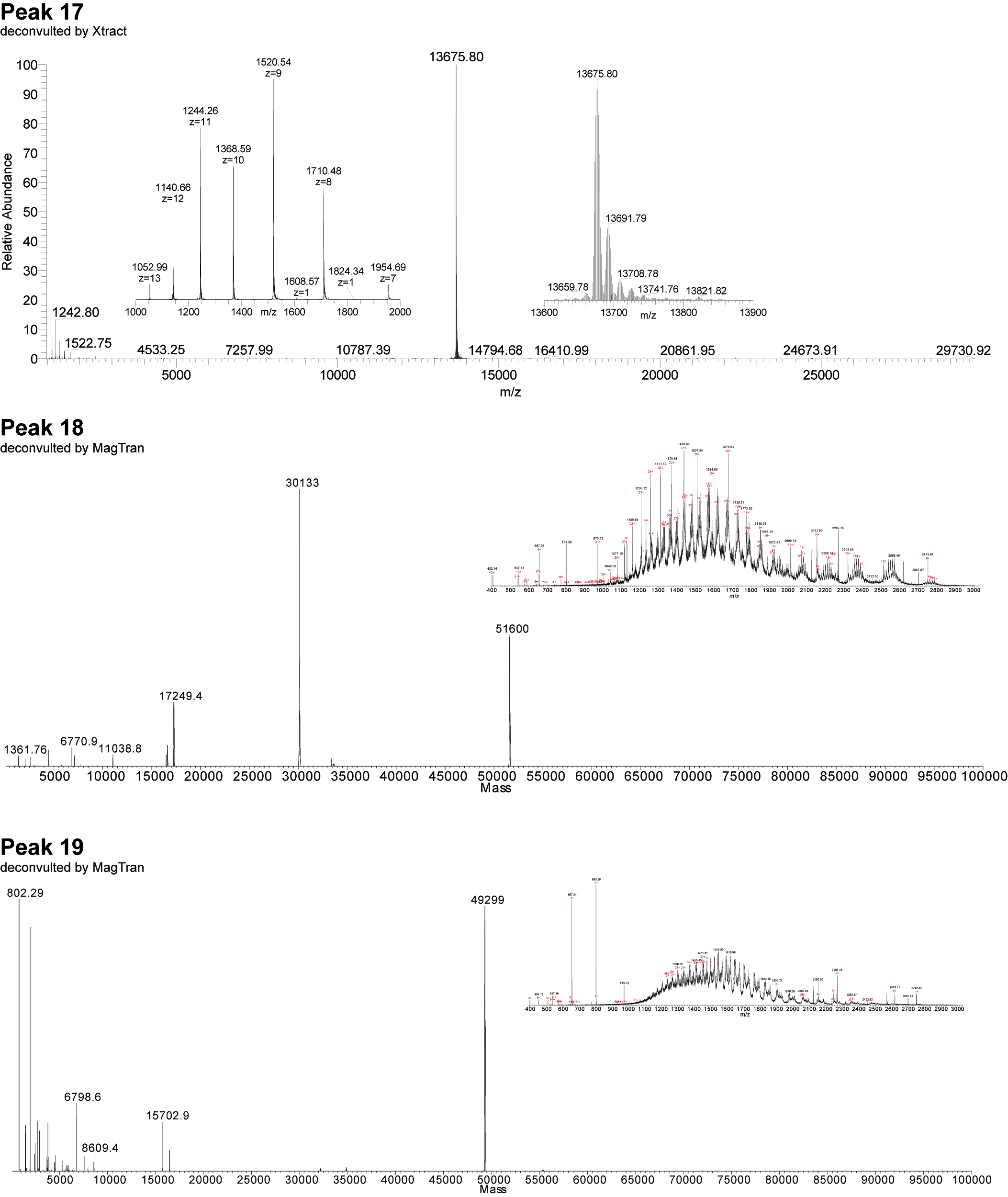

**Supplemental Figure 1. continued**

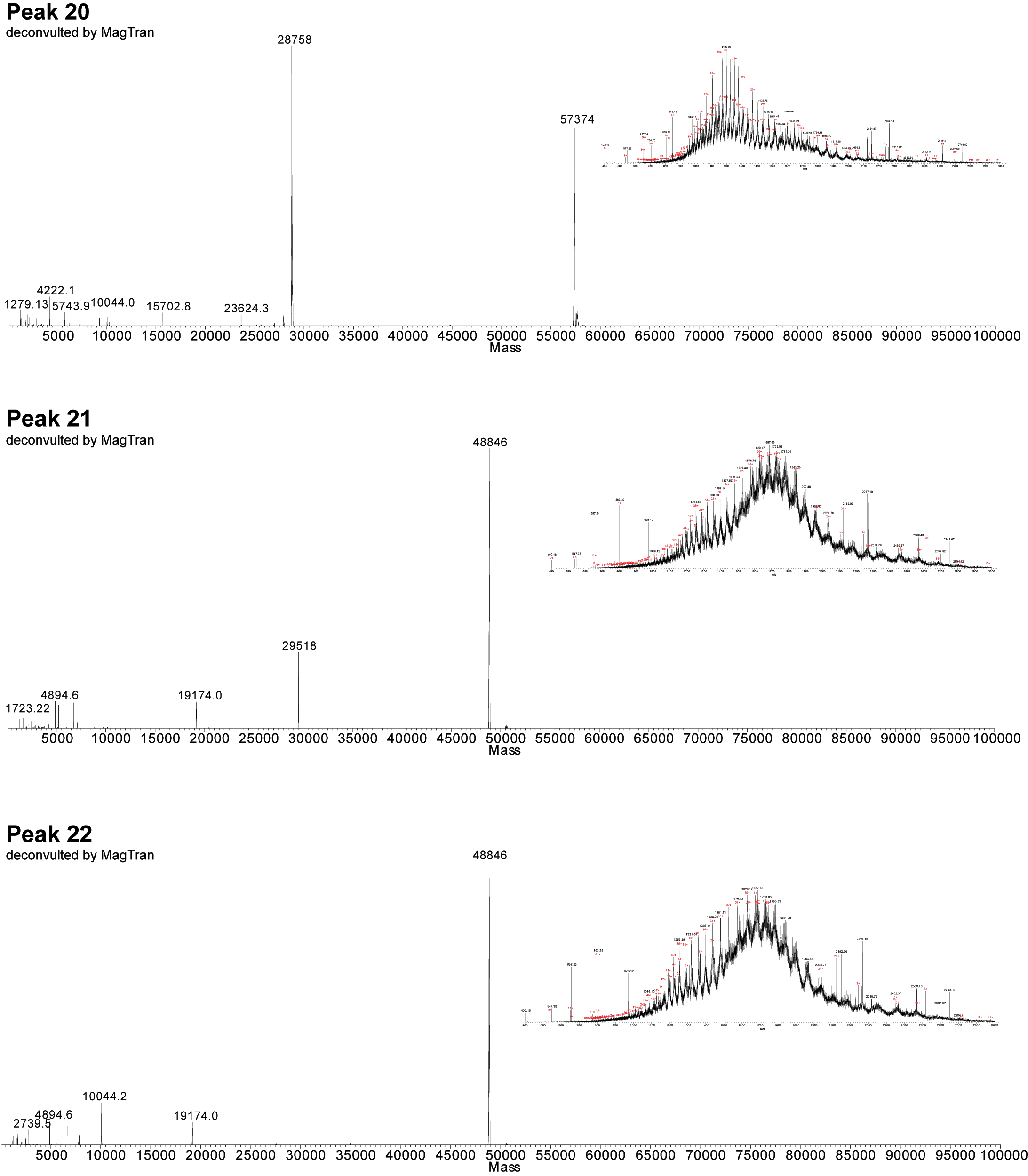

**Supplemental Figure 1. continued**

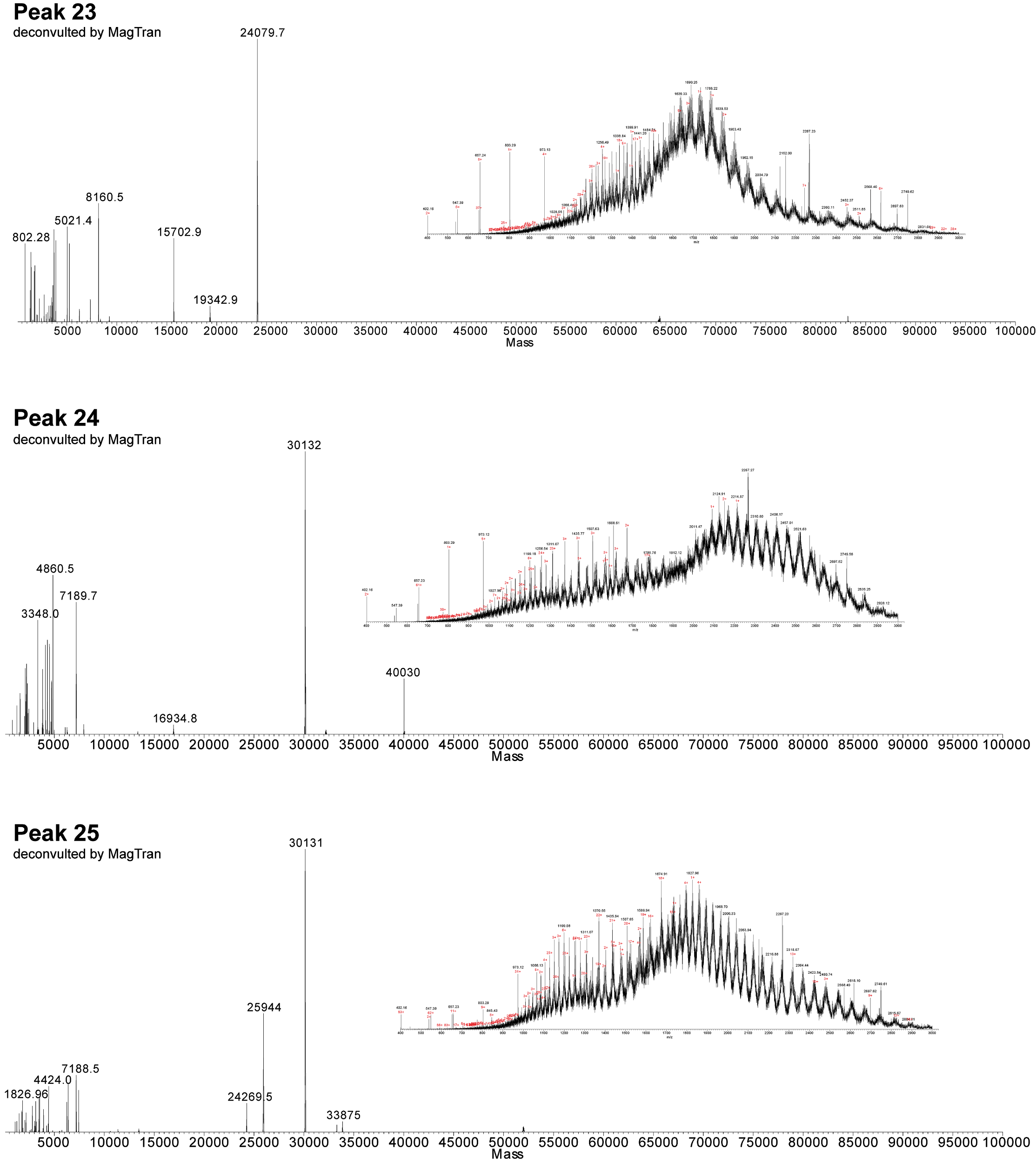

**Supplemental Figure 1. continued**

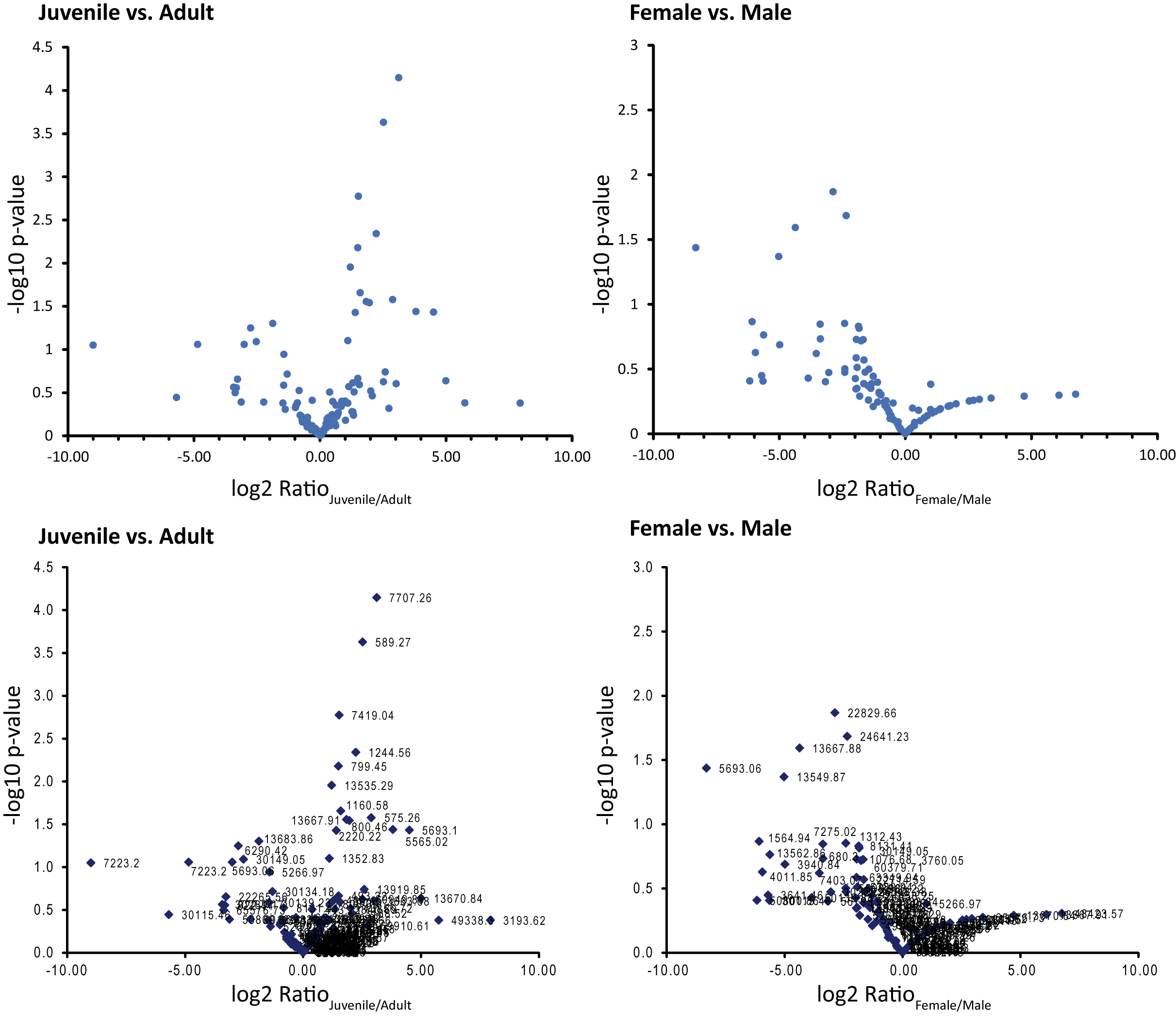

**Supplemental Figure 2.** Vulcano plots of female vs. male individuals and juvenile vs. adult animals. The fold change of proteoform abundance (log2 Ratio) vs. statistical significance (-log10 p-value) is shown. Log2 ratios > 2 or <-2 with -log10 p-values > 1.3 (p-value < 0.05) were considered as significantly differentially expressed proteins.

**Supplemental Table 1. Acute LD50 value of *V. kaznakovi* crude venom.** Determination of LD50 value of V. kaznakovi crude venom in mice following 24h exposure by intraperitoneal injection.

| **Crude venom concentration [mg/kg] (n=5)** | **Dead** | **Live** | **Viability rate [%]** | **Determined LD_50_ value [mg/kg]** |
| --- | --- | --- | --- | --- |
| 5 | 5 | 0 | 0 | **2.59** |
| 2 | 1 | 4 | 80 |  |
| 1 | 0 | 5 | 100 |  |
